# The ROCK1 PHC1 domain interacts with active Rho to transduce cell contraction signals

**DOI:** 10.1101/2025.09.12.675761

**Authors:** Carolin Gierse, Ingrid R. Vetter, Leif Dehmelt

## Abstract

The spatio-temporal regulation of cell contraction is crucial for numerous biological processes. In particular, contraction pulses in the cell cortex contribute to mechanotransduction and tissue rearrangements during embryonic development. We previously identified a signaling network that generates mechanosensitive contraction pulses in adherent mammalian cells via positive and negative feedback regulation of the small GTPase RhoA. Our investigations into the molecular mechanism of this process revealed surprising observations that challenge prevailing models of ROCK1 regulation. In particular, we identified a novel RhoA binding site in the ROCK1 PHC1 tandem domain that is sufficient for dynamic recruitment leading to increased Rho activity within subcellular regions of the cell cortex. AlphaFold-guided mutagenesis supports a direct interaction between these molecules. Functional investigations show that the PHC1 domain is required for efficient recruitment to active Rho, and that it plays a role in the transduction of Rho activity via ROCK1 to Myosin II activation. Based on the newly identified Rho binding site at the C-terminus of ROCK1, we propose a model for ROCK1 activation, which can resolve inconsistencies between previous biochemical and structural studies.

## Introduction

Dynamic changes in cell morphology are essential for many biological processes, such as embryonic development, wound healing or cancer metastasis. These morphological changes are enabled by dynamic rearrangements of the cytoskeleton. In addition to driving cell shape changes, cytoskeletal dynamics are also used by cells to actively probe the mechanical properties of their surroundings (Aguilar-Cuenca et al., 2014). This is achieved by an active probing mechanism, which involves dynamic cell contraction pulses, that were shown to transduce mechanical signals more effectively compared to constant contraction (Cui et al., 2015; Graessl et al., 2017; Nalbant and Dehmelt, 2018). The contractile force in these pulses is generated by the action of non-muscle Myosin II, which shifts anti-parallel actin filaments against each other near the plasma membrane (Niederman and Pollard, 1975; Svitkina et al., 1995).

Pulsatile cell contraction dynamics are a widespread phenomenon, that is observed in various processes that are associated with the function of individual cells and the development of multicellular structures (Martin et al., 2009; Bement et al., 2015; Graessl et al., 2017; Nalbant and Dehmelt, 2018; Kim et al., 2018; Michaux et al., 2018). In several of these systems, pulsatile dynamics were found to be generated by an activator-inhibitor signal network which combines rapid positive and slow negative feedback regulation in the cell cortex near the plasma membrane (Bement et al., 2015; Graessl et al., 2017; Michaux et al., 2018). Detailed analyses in adherent mammalian cells revealed that the active form of the small GTPase Rho recruits its activator GEF-H1 (human ARHGEF2, UNIPROT Q92974) to close a positive feedback loop (Graessl et al., 2017; Kamps et al., 2020), and this fast reaction is coupled to a negative feedback loop, which is mediated by a much slower Myosin-II-dependent process (Lee et al., 2010; Graessl et al., 2017; Kamps et al., 2020) (Figure 4A). The coupling of these processes is dependent on a well-known Rho effector, the Rho kinase ROCK (Figure 4A), via phosphorylation of the Myosin II light chain (MLC) and subsequent activation of Myosin II (Mott et al., 1996; Leung et al., 1996; Amano et al., 1996; Amano et al., 1997; Ishizaki et al., 1997; Graessl et al., 2017).

Earlier studies suggested a relatively simple mechanism for ROCK activation, in which the C-terminal region of ROCK binds and inhibits the N-terminal kinase domain (Chen et al., 2002), and in which this autoinhibition is released by an interaction between active Rho and the ROCK C-terminus (Figure S1A) (Raouf A. Khalil, 2010; Khalil, 2010). However, more recent studies questioned this idea and proposed based on electron microscopy imaging, that ROCK is always in a fully extended conformation, with its long coiled-coil region acting as a spacer to position the kinase domain at a distance of approx. 120 nm from the plasma membrane (Figure S1B) (Truebestein et al., 2015). According to this more recent hypothesis, the N-terminal kinase domain of ROCK would only be able to reach and activate Myosin in the dense cell cortex near the plasma membrane in this fully extended conformation. However, it remains unclear how active Rho would be able to activate ROCK, as all previously described Rho binding sites are located within the coiled-coil domain, far away from the plasma membrane in the fully extended ROCK conformation. Furthermore, these more recent studies failed to confirm the previously proposed interaction between ROCK and active Rho, thus leaving a gap in our understanding, how ROCK and its downstream targets can be activated.

Here, we investigated the interaction between Rho and ROCK1 during pulsatile cell contraction dynamics. We found that full length ROCK1 is quickly recruited to Rho activity pulses within seconds, well before the recruitment and activation of Myosin II reaches its maximum. Surprisingly, this rapid recruitment of ROCK1 was not dependent on any of its previously described Rho interaction domains. Instead, a newly identified interaction surface at the C-terminal PHC1 tandem domain was sufficient for highly effective plasma membrane recruitment to active Rho. Based on predictions using AlphaFold, we identified four amino acids that significantly contribute to this interaction, and we confirmed their importance via mutagenesis. Furthermore, we found that this newly identified interaction contributes to the efficient recruitment of full-length ROCK to the plasma membrane. While our functional investigations confirmed the importance of the PHC1 domain for Myosin activation, the interaction of the PHC1 domain with Rho surprisingly seems to reduce the ability of ROCK1 to activate Myosin. We conclude that the mPHC1 domain contributes to ROCK1 recruitment to active Rho, and that an additional function of this domain apart from localizing to the actin cortex is necessary for ROCK1 to carry out its function to activate Myosin II.

## Results and Discussion

### ROCK1 is dynamically recruited to cell contraction pulses in adherent mammalian cells

In previous studies, we identified and characterized a signaling network in mammalian, adherent cells, which can generate spontaneous cell contraction pulses at the plasma membrane (Figure 1A) (Graessl et al., 2017; Kamps et al., 2020). In these studies, we used total internal reflection fluorescence (TIRF) microscopy to detect the plasma membrane translocation of a fluorescent-labelled Rho effector domain (Benink and Bement, 2005) to measure local Rho activity dynamics within individual pulses (Graessl et al., 2017) and we combined this with measurements of plasma membrane translocation of Rho activating GEFs and of Rho downstream targets, such as F-actin, FHOD1 and Myosin (Graessl et al., 2017; Kamps et al., 2020). These studies revealed that Rho and its activator GEF-H1 can rapidly amplify each other with a minimal temporal time lag of less than 3 seconds, and that the recruitment and activation of downstream targets is significantly delayed (Graessl et al., 2017). In particular, maximal Myosin-II recruitment at the plasma membrane - as detected by TIRF microscopy - occurred about 40s after maximal Rho activation (Graessl et al., 2017). This recruitment was strongly correlated with the formation of centripetal contractile flows, which shows that Myosin II is not only recruited to Rho activity pulses at the plasma membrane but also locally activated at these pulses (Graessl et al., 2017). Here, we used a similar strategy to investigate the dynamics of the Rho effector ROCK within individual cell contraction pulses (Figure 1B-D). Analogous to our previous studies, we simply express a constitutively active, microtubule-binding deficient GEF-H1 C53R mutant to stimulate the generation of local, subcellular activity pulses of the cell contraction signal network, and thereby enhance the sensitivity of our analyses (Krendel et al., 2002; Graessl et al., 2017; Kamps et al., 2020). As shown in middle panels of Figure 1B-D, we confirmed our previous observation (Graessl et al., 2017), that the spatio-temporal plasma membrane recruitment dynamics of fluorescent-labeled GEF-H1 C53R and of the Rho activity sensor are identical. For the following studies, we therefore use GEF-H1 C53R both as a stimulator of cell contraction pulses, and as a sensor to report spatio-temporal Rho activity dynamics.

**Figure 1:**
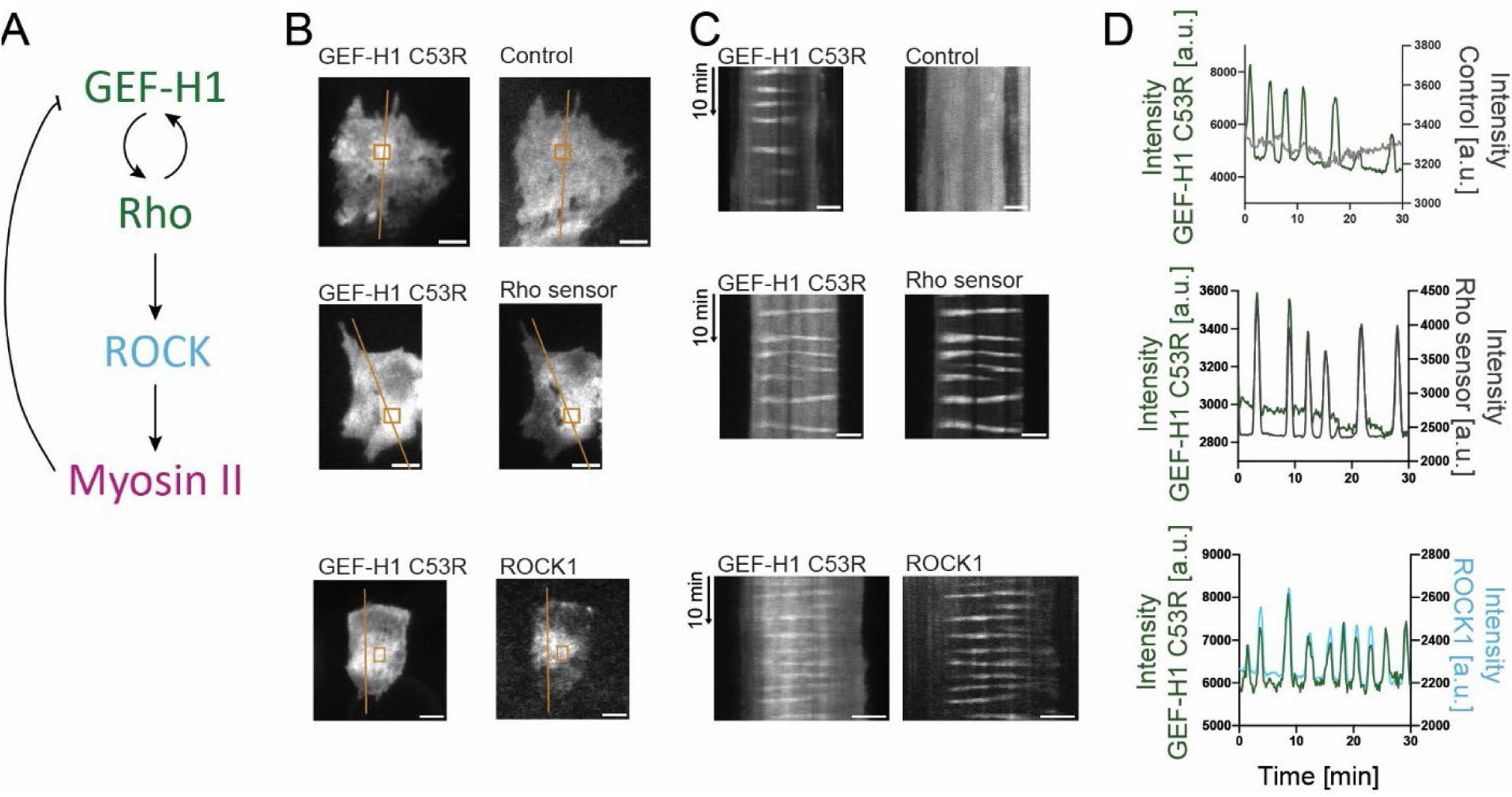
Full length ROCK1 is dynamically recruited to active Rho at the plasma membrane. **A** Topology of the cell contraction signaling network. GEF-H1 activates the small GTPase Rho, which in turn recruits GEF-H1 to the plasma membrane resulting in positive feedback. Rho activates Myosin II via its effector ROCK. Myosin II mediates negative feedback by competing with GEF-H1 (Lee et al., 2010; Graessl et al., 2017; Kamps et al., 2020). **B**-**D** Analysis of ROCK1 plasma membrane recruitment dynamics. **B** TIRF images of representative cells, in which cell contraction network dynamics were stimulated by the expression of the constitutively active GEF-H1 mutant C53R (mApple-GEF-H1 C53R top and middle or mCitrine-GEF-H1 C53R bottom). Cells either co-express delCMV-mCitrine as a volume marker (top), a Rho activity sensor based on the Rhotekin GTPase-binding domain (GBD) (delCMV-mCitrine-Rhotekin-GBD; middle) or fluorescent-labelled full-length ROCK1 (mCherry-ROCK1 full length; bottom). **C** Kymographs corresponding to orange lines in **B**. **D** Kinetics of fluorescence signals in rectangular regions of interest in **B**. Scale bars: 10 µm.

Using this approach, we investigated the spatio-temporal relationship between human full-length mCherry-ROCK1 and its activator Rho (Figure 1B-D, bottom). These experiments clearly show that ROCK1 plasma membrane recruitment closely follows GEF-H1 C53R in space and time throughout the whole cell attachment area with minimal temporal delay. Detailed cross correlation analyses show that ROCK1 is maximally recruited just 3.12 ± 0.56s (mean ± SEM) after GEF-H1 C53R (Figure S1). Analogous experiments with a ROCK2 construct showed similar results (Figure S1D-F).

### Identification of a novel RhoA binding domain that is sufficient for recruitment of ROCK1 to Rho activity pulses

A recent study (Truebestein et al., 2015) has cast doubt on a direct interaction between ROCK1 and active Rho, in particular via the previously characterized Rho binding domain (RBD, residues 994-1009) within its coiled-coil, raising the question how the tight spatio-temporal cross correlation between ROCK1 and GEF-H1 (Figure 1B-D) could be explained (Truebestein et al., 2015). In addition to this canonical RBD region, two additional Rho binding sites were previously identified in ROCK (Figure 2A): The homology region 1 (HR1, residues 420-550) and the Rho-interacting domain (RID, residues 842-931) (Blumenstein and Ahmadian, 2004). These additional potential binding sites are also localized in the coiled-coil domain (Truebestein et al., 2015). At the very C-terminus, ROCK has a Pleckstrin homology (PH) domain with a cysteine-rich (C1) domain inserted within the PH domain (PHC1 tandem domain in the following), and this domain was shown to bind the plasma membrane via a cluster of positive charges (Wen et al., 2008). In other proteins, including Lbc-type GEFs and Raf kinase, PH and C1 domains were found to specifically interact with the active form of small GTPases (Mott et al., 1996; Graessl et al., 2017; Kamps et al., 2020). We therefore hypothesized that this ROCK1 domain might not only interact with the plasma membrane, but might also be able to interact with active Rho.

**Figure 2:**
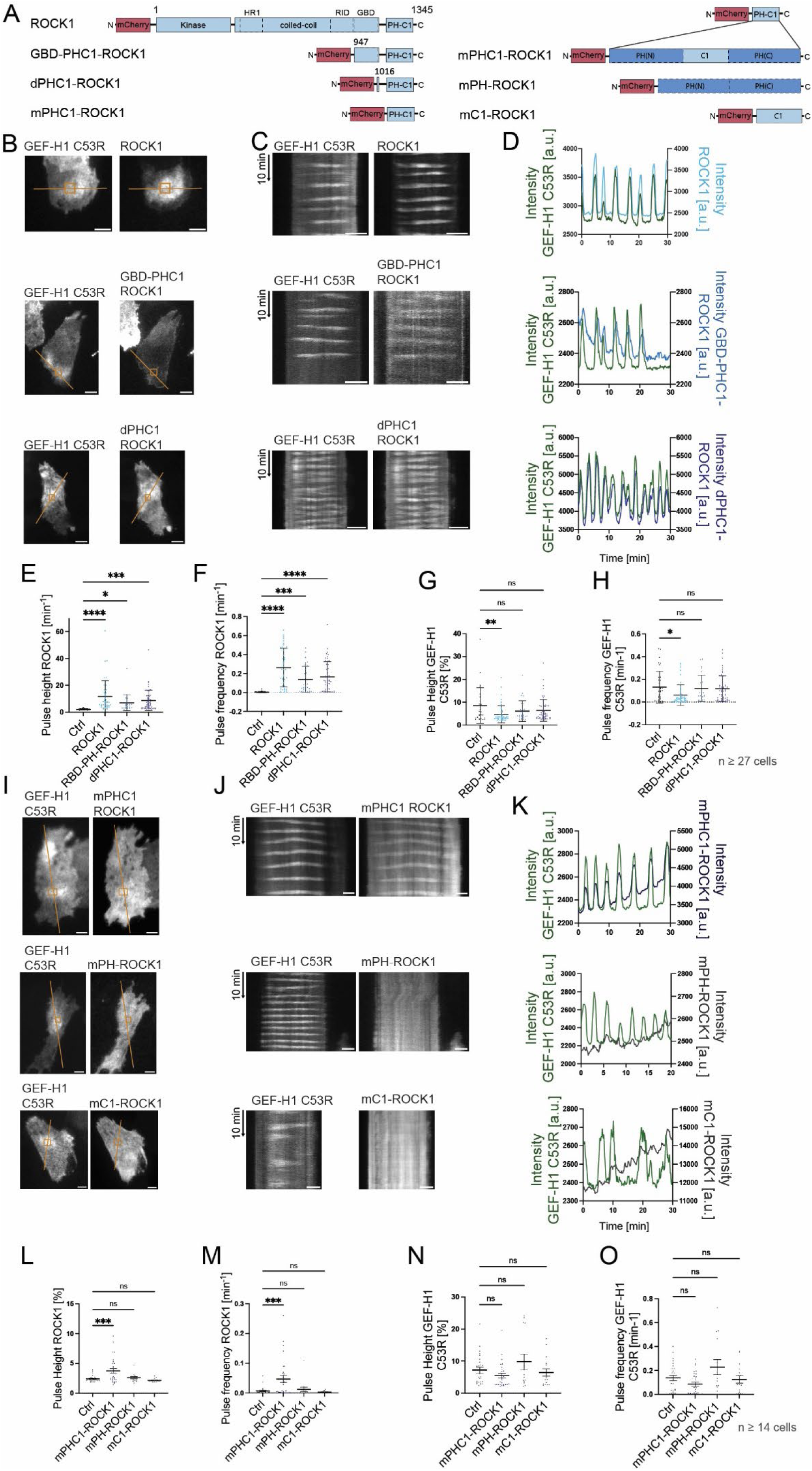
The monomeric PHC1 tandem domain of ROCK1 is sufficient for RhoA activity dependent plasma membrane recruitment. **A** Schematic representation of ROCK1 constructs used in this study. (HR1: homology region 1, RID: Rho-interacting domain, RBD: Rho-binding domain, PH: Pleckstrin homology domain; C1: phorbol esters/diacylglycerol binding domain; the d and m prefix indicate presumed monomeric or dimeric constructs, based on the absence or presence of the coiled-coil region). **B**-**F** Analysis of plasma membrane recruitment dynamics of N-terminal ROCK1 truncation mutants. **B** TIRF images of representative cells co-expressing mCitrine-GEF-H1 C53R and mCherry-ROCK1 full length (top), mCherry-RBD-PHC1-ROCK1 (middle), or mCherry-dPHC1-ROCK1 (bottom). **C** Kymographs corresponding to the orange lines in **B**. **D** Kinetics of fluorescence signals in rectangular regions of interest in **B**. **E-F** Quantitative analysis of plasma membrane recruitment dynamics of mCherry-ROCK1 constructs shown in **B**-**D**, compared to control (empty parental mCherry vector). **G-H** Quantitative analysis of signal network dynamics by measurements of mCitrine-GEF-H1 plasma membrane recruitment in the same experiment as shown in **E-F**. **I-M** Analysis of plasma membrane recruitment dynamics of ROCK1 mPHC domain deletion mutants. **I** TIRF images of representative cells co-expressing mCitrine-GEF-H1 C53R and mCherry-mPHC1-ROCK1 (top), mCherry-mPH-ROCK1 (middle), or mCherry-mC1-ROCK1 (bottom). **J** Kymographs corresponding to the orange lines in **I**. **K** Kinetics of fluorescence signals in rectangular regions of interest in **I**. **L-M** Quantitative analysis of plasma membrane recruitment dynamics of mCherry-ROCK1 constructs shown in **I-K**, compared to control (empty parental mCherry vector). **N-O** Quantitative analysis of signal network dynamics by measurements of mCitrine-GEF-H1 plasma membrane recruitment in the same experiment as shown in **L-M**. Error bars represent standard error of the mean from 3 independent experiments each. *: p < 0.05, **: p < 0.01, ***: p < 0.001, ****: p < 0.0001, scale bars: 10 µm.

To address this question, we generated N-terminally truncated human ROCK1 constructs. The first set of constructs that we investigated either contained both the originally identified canonical RBD (Blumenstein and Ahmadian, 2004) and the PHC1 tandem domain (RBD-PHC1-ROCK1, residues 947-1345) or only the PHC1 tandem domain together with a minimal coiled-coil region to enable its dimerization (dPHC1-ROCK1, residues 1016-1354) (Figure 2A). Thus, RBD-PHC1-ROCK1 only contains the previously characterized canonical RhoA RBD, while dPHC1-ROCK1 does not contain any previously reported RhoA binding site.

We then measured the dynamic recruitment of these constructs to cell contraction pulses in U2OS cells, in which the signal network dynamics were stimulated by GEF-H1 C53R co-expression (Graessl et al., 2017; Kamps et al., 2020) (Figure 2B-D). As expected, RBD-PHC1-ROCK1 plasma membrane recruitment closely correlates with maximal Rho activity indicated by GEF-H1 C53R signals. Interestingly, dPHC1-ROCK1, which contains none of the previously postulated RBDs (Blumenstein and Ahmadian, 2004) equally co-localized with GEF-H1 dynamics. This shows that none of the previously postulated RBDs are required to recruit ROCK1 to sites of increased GEF-H1/Rho activity (Figure 2B-D). Furthermore, quantitative analysis of ROCK1 pulse dynamics (Graessl et al., 2017) confirms that both truncations exhibit pulsatile plasma membrane recruitment dynamics that are statistically indistinguishable (Figure 2E-F).

We next investigated if these ROCK1 constructs affect cell contraction signal network dynamics. To evaluate this, we measured the dynamics of the GEF-H1 signals in the presence of these ROCK1 constructs. These measurements show that ROCK1 wild type had an inhibitory effect, while RBD-PHC1-ROCK and dPHC1-ROCK1 did not influence dynamics, neither concerning GEF-H1 peak height nor GEF-H1 pulse frequency (Figure 2G-H). This was initially surprising, as previous studies suggested an inhibitory, dominant negative effect of the PHC1 domain of ROCK on cell contraction (Leung et al., 1996; Ishizaki et al., 1997; Amano et al., 1997). This shows that the two truncated constructs do not influence oscillatory cell contraction dynamics, and that instead increasing the expression level of the wild-type molecule, which includes the N-terminal kinase domain, inhibits cell contraction signal network dynamics^1^.

To narrow down the minimal binding site for RhoA in ROCK1, we removed the small, dimerizing coiled-coil that was still present in the dPHC1-ROCK1 construct, to obtain a monomeric PHC1 domain (mPHC1, residues 1119-1354, Figure 2A). As shown in Figure 2I-K (top), the plasma membrane recruitment of the mPHC1 domain still clearly and closely correlates with GEF-H1 signals in space and time.

Since both the PH and the C1 domains of other proteins were previously shown to bind small GTPases (Mott et al., 1996; Kamps et al., 2020; Powis et al., 2023), we further dissected the tandem domain to obtain the individual, monomeric PH (mPH-ROCK1, residues 1119-1228 and 1282-1354, connected via a flexible synthetic linker) and C1 (mC1-ROCK1, residues 1229-1281) domains (Figure 2A). The mC1-ROCK1 domain did not exhibit any detectable signal changes in relation to GEF-H1 (Figure 2I-K, bottom), and the mPH-

ROCK1 was only very weakly enriched at local GEF-H1 signals (Figure 2I-K, middle). The automated peak analysis confirmed these observations: mC1-ROCK1 did not show an increase in the ROCK1 pulse height and mPH-ROCK1 showed a very small trend, which was not significantly different compared to the empty control construct (Figure 2L-M). This shows that both domains together are essential for efficient plasma membrane recruitment of the C-terminal region of ROCK1 to active Rho.

In addition, we did not observe any reduction in network activity upon expression of neither mPHC1-, mPH-, nor mC1-ROCK1, further supporting our observation that the PH domain itself has no dominant negative effect on pulsatile cell contraction dynamics (Figure 2N-O).

### Identification of amino acids that are critical for the mPHC1 domain recruitment to Rho activity pulses

To further characterize how mPHC1-ROCK1 is recruited to cell contraction network activity, we employed AlphaFold2 and predicted the potential interaction between this domain and a set of small GTPases. Indeed, an interaction between mPHC1 and GTP bound Rho family GTPases was predicted with high confidence (Figure 3A-B). Compared to GTPases of the Ras, Rap and Rab families, the interface predicted template modeling score (ipTM) for the most studied members of the Rho subfamily, in particular RhoA, RhoB and RhoC, was consistently higher than the 0.8 threshold that is often used to discriminate true vs false positives (Figure S2B). In contrast to other PH or C1 domains, that can selectively bind signaling lipids such as phosphoinositides, or diacylglycerol, respectively, the tandem PHC1 domain of ROCK1 lacks the signatures for binding these molecules (Wen et al., 2008). Instead, the ROCK PHC1 domain has unconventional lipid membrane binding properties that are based on a cluster of positive charges (Wen et al., 2008). Interestingly, these positive charges are located right next to the Rho binding site and oriented in the same direction as the farnesylated, C-terminal Rho plasma membrane anchor (Figure 6E-F and Figure S4C-D). Thus, the positive charges of the PHC1 domain together with the RhoA positive charges and farnesyl moiety might act synergistically to support its ability to bind to the plasma membrane.

**Figure 3:**
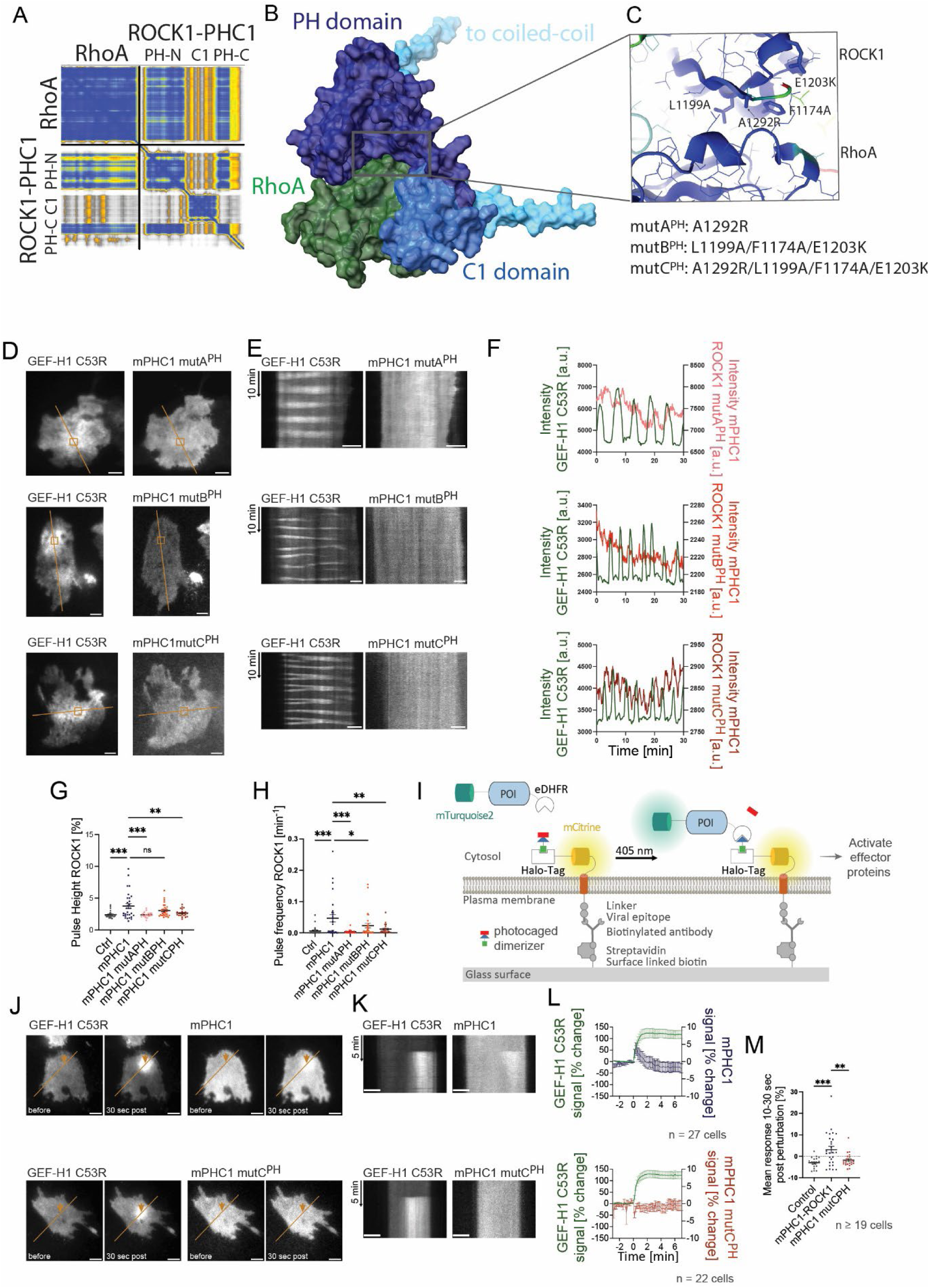
Structural requirements for the interaction between the ROCK1 PHC1-domain and active RhoA. **A**-**C** AlphaFold2 prediction of the PHC1/RhoA complex. **A** Predicted aligned error (PAE) of the predicted PHC1/RhoA complex. **B** Structure of the predicted complex (blue: PHC1-tandem domain; green: RhoA). **C** Magnified view of the predicted binding interface between the PH domain of ROCK1 and RhoA. Amino acids predicted to be most important for the interaction via Protein Interfaces, Surfaces and Assemblies (PISA) are indicated. **D-H** Analysis of plasma membrane recruitment dynamics of ROCK1 mPHC domain point mutants. **D** TIRF images of representative cells co-expressing mCitrine-GEF-H1 and mCherry-mPHC1 mutA^PH^ (top), mCherry-mPHC1 mutB^PH^ (middle) or mCherry-mPHC1 mutC^PH^ (bottom) (**D**). **E** Kymographs corresponding to lines in **D**. **F** Kinetics of fluorescence signals in rectangular regions of interest in **D**. **G-H** Quantitative analysis of plasma membrane recruitment dynamics of mCherry-ROCK1 constructs shown in **D-F**, compared to control (empty parental mCherry vector). **I**-**M** Analysis of plasma membrane recruitment of ROCK1 mPHC domain point mutants to local, light-induced stimulations of the cell contraction signal network. **I** Schematic of the Molecular Activity Painting approach used to introduce local perturbations. In this method, a single pulse of near UV light (405nm) is used to trigger the interaction of eDHFR and a plasma membrane anchored Halo-Tag. This leads to the plasma membrane recruitment and enrichment of a protein of interest (POI) fused to eDHFR. The enriched protein of interest can introduce a local perturbation of signal network activity at the plasma membrane. Total internal reflection fluorescence microscopy (TIRF-M) is used to measure the kinetics of the perturbation and of the downstream response. **J** TIRF images of the mTurquoise2-eDHFR-GEF-H1 C53R perturbation construct (left) and mCherry-mPHC1 (right, up) or mCherry-mPHC1 mutC^PH^ (right, bottom), before and 30 sec after initiation of a perturbation at a local spot of the plasma membrane. Orange arrows indicate the position of photodimerizer uncaging. **K** Kymographs corresponding to lines in **J**. **L** Kinetics of mean fluorescence signals measured at the perturbation spot. **M** Quantitative analysis of the ROCK1 construct response 10-30s after light-induced GEF-H1 perturbation. Error bars represent standard error of the mean (SEM) from 3 independent experiments each. *: p < 0.05, **: p < 0.01, ***: p < 0.001, scale bars: 10 µm.

To validate the AlphaFold2-based prediction experimentally, a Protein Interfaces, Surfaces and Assemblies (PISA) analysis was performed (Collaborative Computational Project, Number 4, 1994), to identify amino acid pairs that contribute most to the predicted interaction between the two proteins. Interestingly, the four amino acids that were predicted to have the strongest effect (A1291, F1174, L1199 and E1203) are highly conserved both in ROCK1 and ROCK2 in vertebrates and invertebrates (Figure S2A). A notable exception is the more distantly related ROCK ortholog let-502 in *C. elegans*, for which none of the four amino acids are conserved. Nevertheless, Alphafold3 predicts a complex between the *C. elegans* ROCK ortholog and either human or *C. elegans* Rho (Figure S5, see Methods for details). Interestingly, the amino acids predicted to be important for these interactions are distinct and additionally involve the C1 domain (Figure S5, R1113 and Y1114 of C1 contact RhoA). In the PH domain, M1029 in *C. elegans* corresponds to L1199 in human and is conserved as a hydrophobic residue. The conformation of the RhoA switch II domain differs in *C. elegans*, which can explain the different conservation of critical amino acid residues. Taken together, these analyses suggest that the predicted Rho binding site in the PHC1 domain of ROCK1 might also contribute to the recruitment of ROCK2 and analogous proteins in other species.

These four amino acids in the PHC1 domain were then mutated to disrupt the predicted interaction with RhoA based on electrostatic and steric properties, with minimal impact on the structure of the individual domains. The following residues were predicted to affect RhoA binding, sorted according to the expected effect: A1291R, F1174A, L1199A and E1203K (Figure 3C; mPHC1 mutA^PH^ : A1291R; mPHC1 mutB^PH^: F1174A, L1199A, E1203K; mPHC1 mutC^PH^: A1291R, F1174A, L1199A, E1203K), and the plasma membrane recruitment of the resulting mutated mPHC1 domains to GEF-H1 activity patterns were investigated.

These experiments showed that two independent sets of mPHC1 domain mutations (the triple mutant mutA^PH^ and the single mutant mutB^PH^), display significantly reduced recruitment to GEF-H1/Rho activity signal dynamics: We measured a strong reduction in the recruitment of a single amino acid mutant (mPHC1 mutA^PH^ : A1291R Figure 3D-F, top), and the independent, non-overlapping triple mutant (mPHC1 mutB^PH^: F1174A, L1199A, E1203K) also showed significantly reduced signals (Figure 3D-F, middle). Combining the triple mutant with the A1291R mutation (mutC^PH^) further reduced the signals (Figure 3D-F, bottom). Quantitative peak analysis confirmed that all three mutants show strongly reduced peak height and frequency compared to the non-mutated mPHC1-ROCK1 domain that served as control (Figure 3G-H).

The peak analysis clearly shows that mPHC1 plasma membrane recruitment coincides with GEF-H1/Rho activity signal network activity, and a more detailed cross correlation analysis confirms a strong spatio-temporal correlation between these signals (Figure S2C). To directly test, if mPHC1 plasma membrane recruitment is indeed caused by increased GEF-H1/Rho activity signal network activity, we employed Molecular Activity Painting, a rapid chemo-optogenetic approach to introduce local perturbations in signal networks, by recruiting a protein of interest to the plasma membrane (Chen et al., 2017; Kamps et al., 2020) (Figure 3I). Here, the constitutively active RhoGEF GEF-H1 C53R is the protein of interest, which is fused to a fluorophore and the heterodimerization domain eDHFR. A Halo-Tag acts as a second heterodimerization domain. This domain is immobilized at the plasma membrane by fusion to an artificial receptor, which is bound to surface-linked antibodies. A photocaged small molecule dimerizer binds covalently to the Halo-Tag, and uncaging by a single pulse of light triggers the plasma membrane recruitment of RhoGEF GEF-H1 C53R-eDHFR. At the plasma membrane, GEF-H1 C53R can activate endogenous Rho (Kamps et al., 2020; Kowalczyk et al., 2022), as well as Rho effectors.

Upon recruitment of GEF-H1 C53R to the plasma membrane, we observed a significant increase in wildtype mPHC1-ROCK signals. This confirms a causal link between GEF-H1-induced cell contraction signal network activity and mPHC1-ROCK1 plasma membrane recruitment (Figure 3J-K). Furthermore, we carried out the exact same experiment with the quadruple mPHC1 mutant (mutC^PH^) and did not detect any local enrichment of this construct (Figure 3J-K).

Interestingly, the wildtype mPHC1 response to the GEF-H1 C53R perturbation is only transient (Figure 3L). This further strengthens the hypothesis that the PHC1 domain of ROCK1 directly interacts with Rho since Rho activity shows the same transient behavior upon recruitment of GEF-H1 C53R (Kamps et al., 2020; Kowalczyk et al., 2022).

### Both the PHC1 domain and the RBD of ROCK1 contribute to its recruitment to active Rho/GEF-H1

To further investigate the functional relevance of the PHC1 domain in ROCK1, we dissected its potential roles in interacting with active Rho at the plasma membrane. First, we introduced the four point mutations into the PHC1 domain of human full-length ROCK1 to examine the role of the PHC1/Rho interaction in the recruitment to cell contraction pulses. As shown in Figure 4A-C (bottom), we found that ROCK1 full length mutC^PH^ still exhibits pulsatile behaviour that closely correlates with GEF-H1 in space and time, presumably due to the presence of the other, previously identified Rho binding sites (Leung et al., 1996; Fujisawa et al., 1996; Blumenstein and Ahmadian, 2004; Dvorsky et al., 2004; Tu et al., 2011). This correlation was similar compared to wild-type ROCK1 (Figure 4A-C top). However, analysis of the peak height showed a trend that this mutant was less effectively recruited to GEF-H1 at the plasma membrane (p=0.3791,Figure 4D). Notably, the more sensitive measurement of the standard deviation of fluorescence intensity signals showed significantly reduced dynamics of the ROCK1 full length mutC^PH^ compared to non-mutated ROCK1 full length at the plasma membrane, further supporting the idea that the newly identified RhoA binding domain contributes to ROCK1 recruitment (Figure 4E).

**Figure 4:**
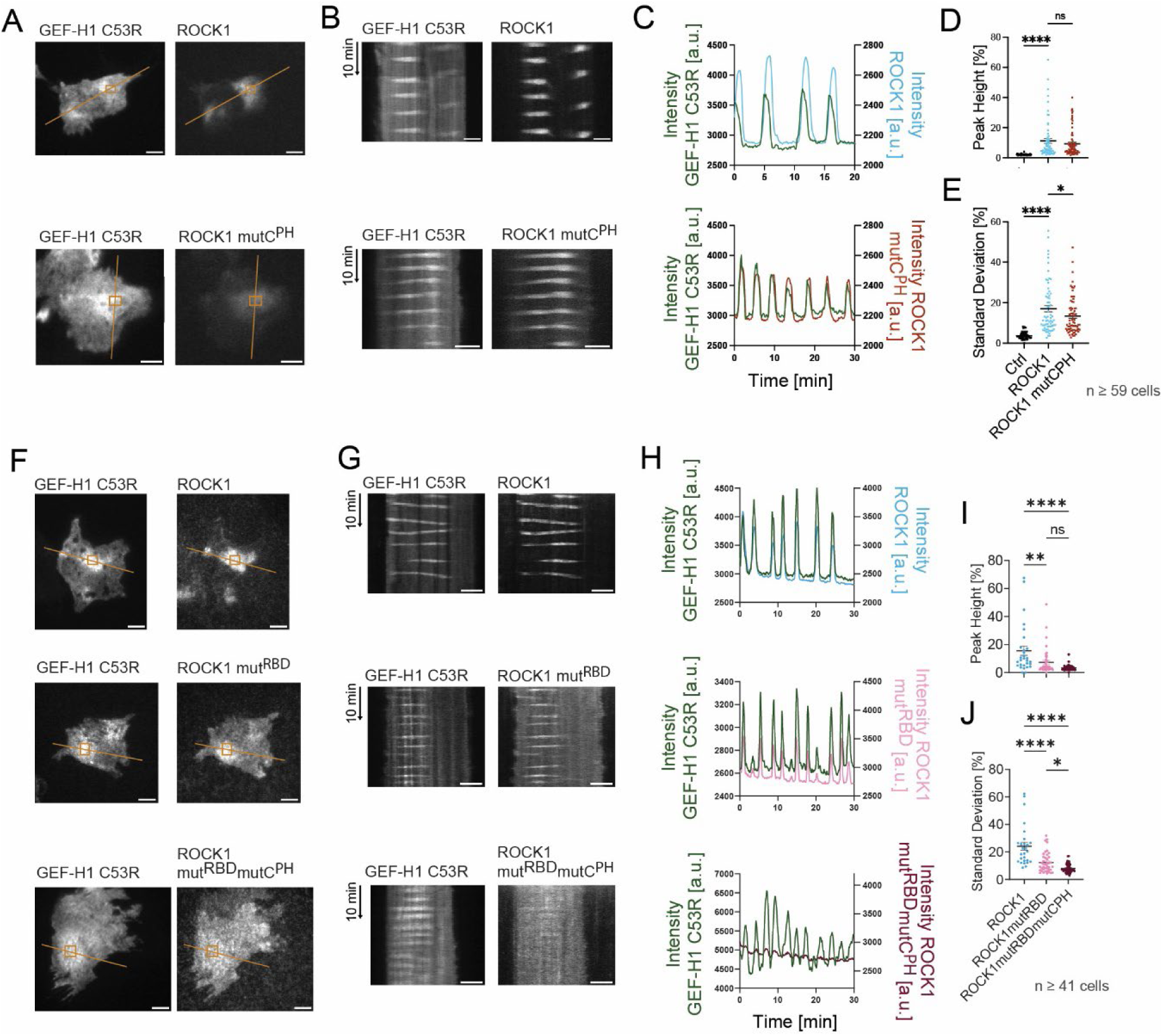
Functional role of the PHC1 domain in the plasma membrane recruitment of ROCK1 to Rho activity dynamics. **A**-**E** Analysis of plasma membrane recruitment dynamics of ROCK1 full length point mutants. **A** TIRF images of representative cells co-expressing mCitrine-GEF-H1 and mCherry-ROCK1 (top) or mCherry-ROCK1 mutC^PH^ (bottom) (A). **B** Kymographs corresponding to lines in **A**. **C** Kinetics of fluorescence signals in rectangular regions of interest in **A**. **D**-**E** Quantitative analysis of plasma membrane recruitment dynamics of mCherry-ROCK1 signals shown in **A**-**C**. **D** Quantification of pulse height (ROCK1 vs. ROCK1mutC^PH^ p=0.3791). **E** Quantification of the local standard deviation over time (ROCK1 vs. ROCK1mutC^PH^: p=0.0465; 4 independent experiments). An empty parental mCherry construct served as control in **D**-**E**. **F-J** Analysis of plasma membrane recruitment dynamics of ROCK1 full length point mutants. **F** TIRF images of representative cells co-expressing mCitrine-GEF-H1 and mCherry-ROCK1 (top), mCherry-ROCK1 mut^RBD^ (middle), or mCherry-ROCK1 mut^RBD^ mutC^PH^ (bottom). **G** Kymographs corresponding to lines in **F**. **H** Kinetics of fluorescence signals in rectangular regions of interest in **F**. **I-J** Quantitative analysis of plasma membrane recruitment dynamics of mCherry-ROCK1 signals shown in **F-H**. **I** Quantification of pulse height. **J** Quantification of the local standard deviation over time. (3 independent experiments). *: p < 0.05, **: p < 0.01, ****: p < 0.0001, scale bars: 10 µm.

To further characterize the recruitment of ROCK1 to cell contraction signal network activity, we also specifically investigated the previously identified RBD region within the coiled-coil region of ROCK1. In previous work, mutation of N1004T and L1005T was shown to strongly suppress the interaction of ROCK1 with active RhoA (Leung et al., 1996). We therefore investigated how these mutations affect the plasma membrane recruitment of ROCK1 to contraction signal network pulses, analogous to the PH domain point mutations described above. As shown in Figure S3, the isolated RBD region itself is recruited to GEF-H1 C53R pulses, however, to a much lesser degree compared to full length ROCK1. The N1004T and L1005T mutations completely abolished this recruitment, showing that these residues are indeed critical for the interaction of the RBD region with active Rho. Introducing these mutations in full length ROCK1 (ROCK1mut^RBD^;Figure 4F-H, middle) significantly reduced recruitment to GEF-H1 C53R pulses, however, dynamics were still clearly detectable (Figure 4F-H (middle) and I-J). Introducing both the mutC^PH^ and the mut^RBD^ mutations (ROCK1mut^RBD^mutC^PH^) nearly abolished ROCK1 plasma membrane recruitment (Figure 4F-H (bottom) and I-J). In particular, the more sensitive measurement of recruitment dynamics via the standard deviation of fluorescence intensity signals was significantly reduced for ROCK1mut^RBD^mutC^PH^ compared to ROCK1mut^RBD^. Taken together, we conclude that the RBD and the PHC1 domains act synergistically to target ROCK1 to the plasma membrane.

### The PHC1 domain of ROCK is critical for the transduction of Rho signals to activate Myosin

We next asked the question, if ROCK1 activity is exclusively regulated by plasma membrane targeting, or if other features of Rho/PHC1 domain interaction also play a role. To independently dissect these two features of ROCK1, we again employed the Molecular Activity Painting assay (Figure 3I) (Chen et al., 2017). Using this method, we first recruited non-mutated, full length ROCK1 to the plasma membrane independently of its endogenous targeting mechanism via photochemical dimerization. This artificial enrichment of ROCK1 lead to a significant increase in fluorescent-labelled Myosin IIa signals at the plasma membrane, indicating that increased local ROCK1 concentrations are sufficient to increase local ROCK1 signaling towards Myosin (Figure 5A-C, ROCK1). Furthermore, the local Myosin IIa enrichment was associated with a strong stimulation of contractile Myosin flows, showing that Myosin IIa is not only recruited but also activated by ROCK1 (Figure5B, top and Figure 5D, top).

**Figure 5:**
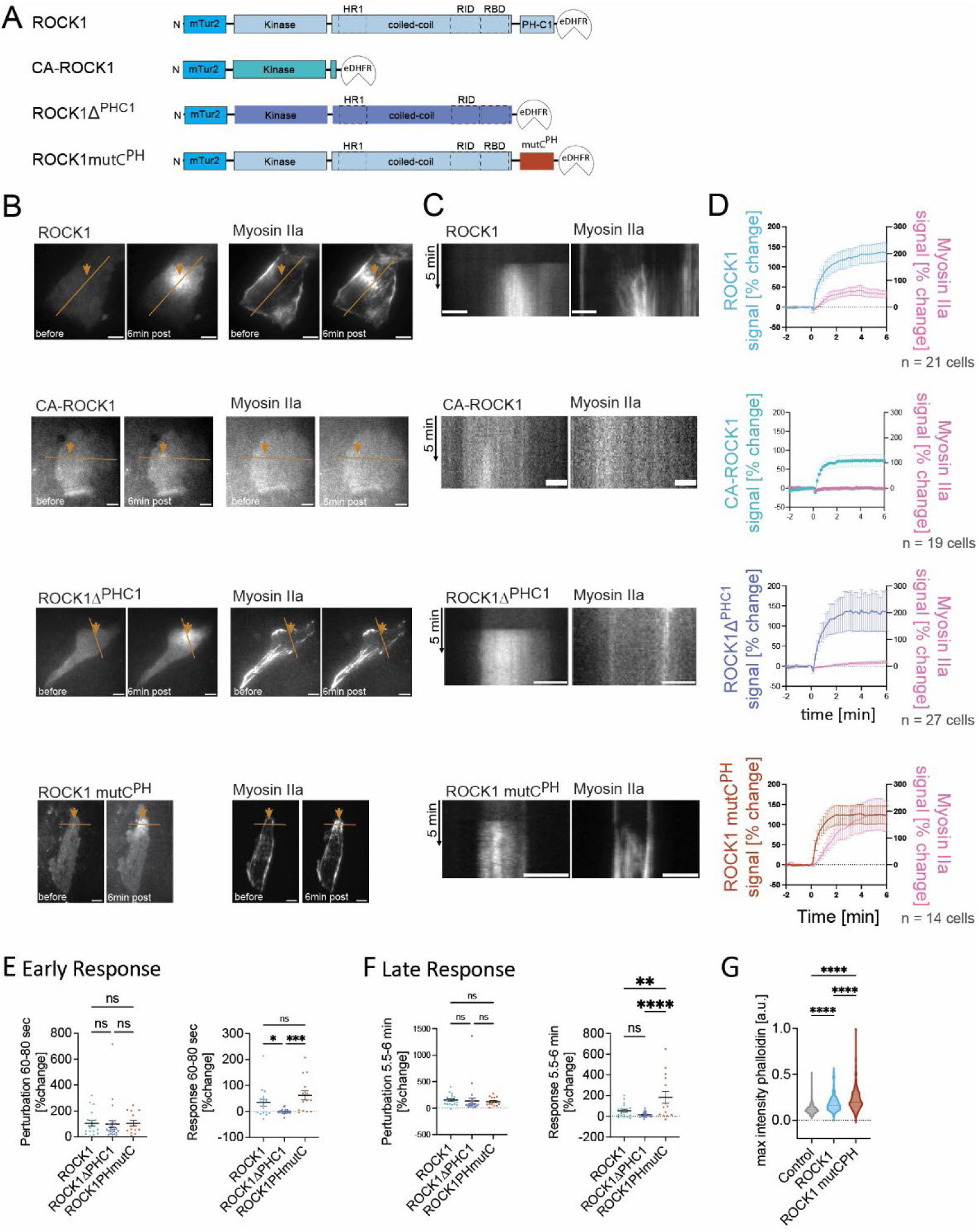
Functional role of the ROCK1 PHC1 domain in the transduction of Rho signals to activate Myosin II. **A** Schematic representation of ROCK1 constructs used in this figure. (HR1: homology region 1, RID: Rho-interacting domain, RBD: Rho-binding domain, PH: pleckstrin homology domain; C1: phorbol esters/diacylglycerol binding domain; mTur2: mTurqoise2; eDHFR: *E. coli* dihydrofolate reductase) **B**-**F** Analysis of plasma membrane recruitment of ROCK1 constructs to local, light-induced stimulations of the cell contraction signal network. **B** TIRF images of the mTurquoise2-ROCK1 (top, left), CA-ROCK1 (middle1, left) ROCK1Δ^PHC1^ (middle2, left), or ROCK1 mutC^PH^ (bottom, left) perturbation constructs and of the mCherry-Myosin IIa response (right), before and 6 min after initiation of the light-induced perturbation. Orange arrowheads indicate the position of photodimerizer uncaging. **C** Kymographs corresponding to lines in **B**. **D** Kinetics of mean fluorescence signals measured at the perturbation spot. **E** Mean perturbation intensity (left) and mean response (right) from 60-80 seconds post perturbation. **F** Mean perturbation intensity (left) and mean response (right) from 5.5-6 min post perturbation. **G** Maximal intensity of Atto488 phalloidin staining of filamentous actin (F-Actin) in fixed cells treated with either ROCK1 (blue), ROCK1 mutC^PH^ (red), or the empty parental construct as control (gray). 3 independent experiments each. *: p < 0.05, **: p < 0.01, ***: p < 0.001, ****: p < 0.0001, scale bars: 10 µm.

We then tested if the ROCK1 kinase-domain by itself is sufficient to stimulate Myosin activity at the plasma membrane. We based this perturbation construct on the previously established CA-ROCK1 mutant (Ishizaki et al., 1997), which only includes the kinase domain and a small region of the coiled-coil to enable dimerization, and was shown to be constitutively active in cells (Figure 5A, CA-ROCK1). Strikingly, enriching this constitutively active mutant at the plasma membrane did not lead to any significant recruitment of Myosin, showing that regions of ROCK1 downstream of the kinase domain, within the coiled-coil and/or the PHC1 domains are essential for its ability to activate Myosin (Figure 5B-C, CA-ROCK1).

Next, we specifically deleted the PHC1 tandem domain of ROCK1 to evaluate its role in Myosin activation. As demonstrated in Figure 5B-C (ROCK1Δ^PHC1^), ROCK1 lacking the PHC1 domain is severely limited in its ability to activate Myosin. Although the artificial recruitment of ROCK1 and ROCK1Δ^PHC1^ to the plasma membrane is equal, a strongly reduced co-recruitment of Myosin at the membrane is observed for the ROCK1Δ^PHC1^ mutant. This shows that the PHC1 domain is critical for the ability of plasma membrane localized ROCK1 to activate Myosin (Figure 5E-F).

We next investigated the role of the Rho/PHC1 domain interaction for Myosin activation by introducing the mutC^PH^ mutations into the perturbation construct. Interestingly, we found that plasma membrane-targeting of ROCK1 full length mutC^PH^ induced an even stronger Myosin co-recruitment compared to ROCK1 full length wildtype (Figure 5B-C and E-F, ROCK1 mutC^PH^), which was again associated with strong contractile Myosin flows (Figure 5B, bottom and Figure 5D, bottom). Although the artificial plasma membrane recruitment of the mutated and non-mutated forms of full length ROCK1 were with similar efficiency, we measured a more than 3 times higher amount of Myosin IIa in the presence of these mutations. Further examination of the kinetics of Myosin recruitment to ROCK1 perturbations reveals that there is no significant difference between ROCK1 and ROCK1 mutC^PH^ during the initial phase of recruitment within the first 60s after perturbation (Figure 5D). In contrast, the augmented Myosin activation only gets apparent at later timepoints within the following minutes (Figure 5E). These observations therefore suggest that the interaction between active Rho and the ROCK1-PHC1 domain has an inhibitory effect on the ROCK1 kinase domain that acts with a delay of about 1 minute.

We next wondered, if this inhibitory effect is dependent on the enforced plasma membrane localization of ROCK1 by the molecular activity painting method and measured the response of long-term overexpression of ROCK1 full length constructs either in the presence or absence of the mutations in the PHC1 domain. In support of an inhibitory function of the Rho PHC1 domain interaction, we found that overexpression of ROCK1 full length mutC^PH^ stimulated the formation of strongly stained F-actin structures, which are typical for acto-myosin-based contractile stress fibers (Figure 5F). In contrast, non-mutated ROCK1 full length was much less effective in stimulating such contractile F-actin structures. This observation was particularly striking considering the reduced recruitment of ROCK1 full length mutC^PH^ to patterns of GEF-H1/Rho activity signals (Figure 4A-F).

Taken together, our experiments show, that the PHC1 tandem domain is on the one hand required to transduce ROCK1 activity to activate Myosin II (Figure 5B-C, ROCK1Δ^PHC1^) and that its interaction with active Rho has a delayed inhibitory effect that limits this transduction (Figure 5E).

## Discussion

Our observations clarify several important questions concerning the mechanism, how ROCK can transduce signals from activated, GTP-bound Rho GTPases to activate Myosin II. First, we show that ROCK1 kinase is strongly and very quickly recruited to local pulses of Rho activity at the plasma membrane with minimal temporal delay. We consider this recruitment as an important first step for the spatio-temporal regulation of ROCK1 in cells. Furthermore, we identified a new Rho binding site at the very C-terminus of ROCK1, within its PHC1 domain, which is sufficient for recruitment to local Rho activity pulses.

This new interaction domain can reconcile a central discrepancy in a recent model for the regulation of ROCK: The “molecular ruler” hypothesis proposed by Truebestein et al, in which the approx. 90-100nm long coiled-coil of ROCK1 is always in a fully elongated conformation to position the kinase domain in sufficient distance from the plasma membrane and to reach Myosin II motors in the dense and crowded cell cortex (Truebestein et al., 2015; Chugh and Paluch, 2018). However, in this fully elongated conformation, the plasma membrane-bound ROCK1 activator Rho is not able to reach the previously known binding site within the ROCK1 coiled-coil (RBD, residues 947-1345, Figure 6A). In contrast, the new Rho binding site in the ROCK1 PHC1, that we now report in our study, can interact with farnesylated Rho at the plasma membrane even when ROCK1 is present in its fully extended conformation. Thus, the Rho-activity dependent recruitment of ROCK1 can thereby link upstream Rho activity at the plasma membrane to kinase activity at a suitable distance to bridge the dense cell cortex for downstream Myosin activation (Figure 6B).

**Figure 6:**
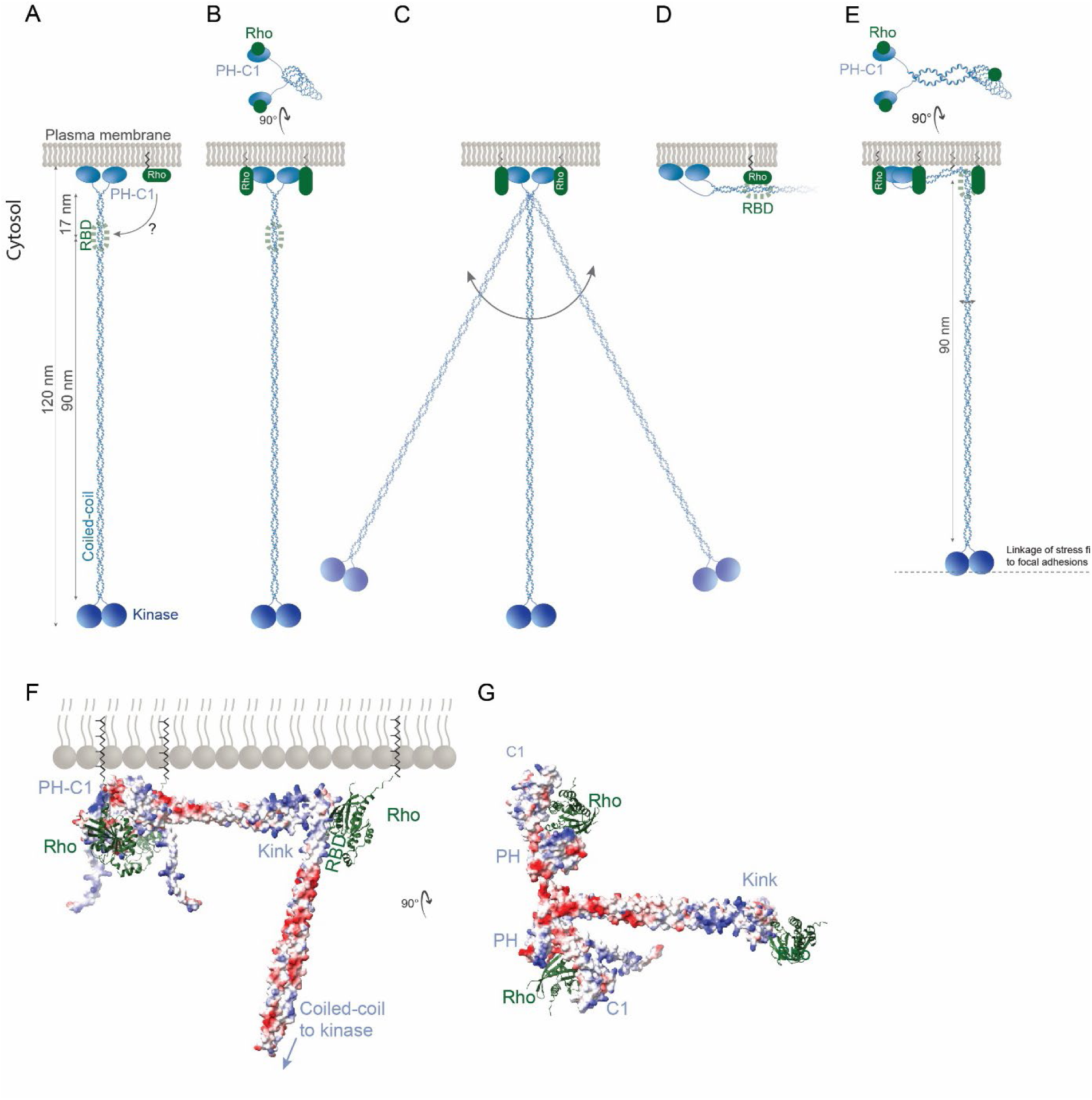
Proposed mechanism for the transduction of Rho signals to myosin II by ROCK1. **A** Schematic of “Molecular Ruler Model” (Truebestein et al., 2015). Based on this model, ROCK binds constitutively to the plasma membrane via the PH domain in its fully extended conformation. However, it is unclear how ROCK can be regulated in this model, as the previously identified ROCK1 RBD region is too far away from active Rho in the plasma membrane. **B** The newly identified Rho interaction with the ROCK1 PHC1 tandem domain can reconcile this discrepancy of the “Molecular Ruler Model”. **C** However, since the linkers that connect the Rho-bound PHC1 domains of ROCK to the coiled-coil domain are flexible, ROCK can pivot back and forth in its extended conformation. This could effectively reduce the average distance between the ROCK1 kinase domain and the plasma membrane and thereby reduce its efficiency to activate Myosin within the dense cell cortex. **D** Binding of ROCK1 to plasma membrane bound Rho via its canonical RBD would stabilize an orientation of the fully-extended coiled-coil parallel to the plasma membrane, reducing the distance between the kinase domain and the plasma membrane to a minimum. In this conformation, ROCK would not be able to reach Myosin within the dense cell cortex. **E** A more stable, perpendicular orientation of ROCK1 could be realized by simultaneous binding of up to 3 plasma membrane-bound Rho molecules in a ROCK1 conformation that includes a predicted kink in the coiled-coil structure (see also Figure S4A-B). **F** Structural model of 3 RhoA-GTP molecules bound to one ROCK1 dimer. The electrostatic surface of the ROCK1 dimer is shown together with the RhoA molecules depicted in ribbon representation. In the presence of the kink in the coiled-coil, two RhoA molecules can bind to the two PHC1 domains, and an additional RhoA molecule can bind to one side of the RBD region. In this configuration, ROCK1 can interact with the plasma membrane via its positively charged surfaces in the PHC1 domains and C-terminal to the RBD region and via its interactions with multiple plasma membrane bound RhoA molecules (see Methods for details). **G** Top view of the structural model shown in **F**.

However, our studies showed that the previously identified RBD region in the coiled-coil plays an important role as well and contributes significantly to ROCK1 recruitment to the plasma membrane. This raises the question, why so many potential Rho interaction domains exist in ROCK. Conceptually, while the PHC1 domains are sufficient to target ROCK1 to the plasma membrane, their flexible linkage to the coiled-coil domain cannot stabilize the perpendicular orientation of ROCK1. Thus, in this state, ROCK might still be able to pivot back and forth at the plasma membrane and might therefore not be able to effectively reach Myosin II within the dense actomyosin cortex (Figure 6C) (Chugh and Paluch, 2018). The canonical RBD region in the coiled-coil is even less suitable to stabilize a perpendicular orientation of the fully-extended coiled-coil. If for example, only one RBD would bind to plasma membrane bound Rho (Truebestein et al., 2015), the whole extended coiled-coil domain would be oriented parallel to the plasma membrane and therefore would not be able to bridge the distance to reach Myosin in the dense cell cortex (Figure 6D).

Intriguingly, based on the well-known heptad and hendecad repeat sequence motifs of coiled-coils (McLachlan and Stewart, 1975; Stetefeld et al., 2000; Truebestein and Leonard, 2016) and on Alphafold2 predictions, the coiled-coil domain is expected to form a kink directly C-terminal to the RBD region (Figure S3A-B). This kink would enable flexibility in the angle between the upstream and downstream coiled-coil structures and would enable a conformation, in which at least one Rho molecule at the plasma membrane can bind the RBD region in the coiled coil (Figure 6E-G). Together with up to two additional Rho molecules that are sterically able to bind two PHC1 domains, this could lead to a much more stable association of ROCK1 with the plasma membrane, which is based on up to three anchor points (Figure 6E-G; compare to loose linkage in Figure 6B-C). Furthermore, clusters of positive charge at the base of the PHC1 domains (Wen et al., 2008), at the kink near the RBD region (AlphaFold prediction, Figure6F-G), and in the helices immediately downstream of this region (Figure 6F-G) might also stabilize the plasma membrane association of this domain and might be able to facilitate a perpendicular orientation of the upstream coiled-coil domain (Figure 6E). Based on this idea, the perpendicular orientation of ROCK1 that is required to efficiently transduce Rho activity to activate Myosin would rely on both the PHC1 domain and the RBD region.

Our findings raise the question if the recruitment of ROCK1 to the plasma membrane in an extended conformation is sufficient to activate Myosin II, or if additional mechanisms also play a role. Strikingly, we found that the artificial targeting of full length ROCK1 via a photochemical dimerization system to a small, photoactivated region of the plasma membrane is indeed sufficient to strongly and highly locally activate Myosin IIa in cells, resulting in a substantial, local cell contraction response. This activation was dependent on several properties of ROCK1. First, artificial plasma membrane targeting of a construct that was missing the majority of the coiled-coil was not able to induce any detectable Myosin activation, showing that this part of ROCK1 is essential, and further supporting the idea of the molecular ruler hypothesis (Figure 5B-C, CA-ROCK1). In addition, we found that the PHC1 domain also plays an important functional role, even if ROCK1 with an intact coiled-coil is efficiently targeted to the plasma membrane via the photochemical dimerization system. The three anchor point structure proposed above (Figure 6E-G) might serve as an explanation for this observation: In the absence of the PHC1 domains, only one plasma membrane associated Rho molecule can bind the RBD region in the coiled coil, which might not be stable enough for a sustained, perpendicular orientation of the coiled-coil and might thereby not be able to efficiently activate Myosin. In the presence of the PHC1 domains, up to three plasma membrane anchor points could be engaged by the ROCK1 C-terminus. It should be noted here, that these anchor points might not require the continued presence of plasma membrane associated Rho, but could also at least partially be facilitated by the strong positive charges in the PHC1 domains, the kink region and the helices that connect these regions.

One of the most surprising results of our studies was that the interaction of the mPHC1 domain with Rho can also reduce the ability of ROCK1 to activate Myosin. Interestingly, this decrease was only observed after a short delay of about 1 minute (Figure 5 A-C, bottom), while ROCK1 recruitment was still increasing. This delayed reduction of ROCK1 activity indicates the presence of a negative feedback regulation. Such negative feedback could for example be part of a failsafe mechanism that might be able to prevent excessive Myosin activity that could potentially be damaging to cells, in particular within tissues in which cells are mechanically coupled to each other. Our earlier work already revealed a much more effective failsafe mechanism that can shut down Myosin activity to baseline within 2 minutes via a negative feedback loop that involves GEF-H1 and Rho activity (Graessl et al., 2017)^2^. Here, our investigations indicate that an additional failsafe might also operate between Myosin and ROCK1, in particular in a situation, in which the regulation of ROCK1 plasma membrane targeting is lost due to the enforced plasma membrane targeting via the photochemical dimerization system. It is currently unclear, how this feedback mechanism might operate. One possibility is that the mPHC1 domains compete with other Rho effector domains, like for example the RBD in the coiled coil region, or effectors that mediate Rho activity amplification (Graessl et al., 2017), and thereby reduce the transduction of endogenous Rho activity towards Myosin at the later timepoints after inducing the ROCK perturbation. In this context, Rho activity amplification might be particularly critical. This amplification is mediated by a positive feedback loop, in which active Rho recruits its activator GEF-H1 (Graessl et al., 2017; Kamps et al., 2020), via a binding site that is completely blocked by the Rho-PHC1 interaction (see crystal structure PDB entry 8BNT). Thus, the PHC1 domains could impede this activity amplification by simultaneously binding and sequestering active Rho, and thereby reducing its ability to further recruit GEF-H1. This sequestration would thereby limit positive feedback amplification of Rho and thus avert potentially excessive Myosin activation in the cell.

In conclusion, we revealed a new Rho interacting domain in the PHC1 domain at the extreme C-terminus of ROCK1 that is sufficient to locally enrich this central signal molecule in subcellular regions of the plasma membrane. Our functional characterization supports an extension to the molecular ruler model, in which several regions of ROCK1, including the coiled-coil and the PHC1 domain, are essential to transduce Rho activity at the plasma membrane to the distant kinase domain of ROCK1.

## Materials and Methods

### Cell Culture

U2OS cells were cultured in DMEM + GlutaMAX^TM^ gibco Ref.: 31966-047 (10% FBS gibco Ref.: A5256701, 10 U/ml Penicillin + 100 µg/ml Streptomycin PAN Biotech Cat.No.: P06-07100). The cells were maintained using standard culture techniques (Trypsin/EDTA 0.05%/0.02% PAN Biotech Cat.no.: P10-02355P, TC dishes Cell+ Sarstedt) at 37 °C and 5% CO-. For standard live cell imaging, cells were plated onto 10 µg/ml collagen-I coated LabTek glass surface slides (Thermo Fischer Scientific) one day prior transfection. All plasmids were transfected into U2OS cells using Lipofectamine^TM^ 2000 (ThermoFisher scientific, Cat.no.: 11668019).

### Plasmid construction

The expression construct CMV-mCitrine-GEF-H1 C53R^3^ coding for mutated GEF-H1 (*Arhgef11,* human), was generated via a Gibson assembly. Briefly, EGFP-GEF-H1 C53R (Krendel et al., 2002) was utilized as the backbone after restriction digestion using EcoRI and KpnI, and delCMV-mCitrine (Nanda et al., 2023) was used as a template to amplify the mCitrine coding sequence using 5’-ctcgtttagtgaaccgtcagaattcgccaccatggtgagcaagggcgag-3’ as forward and 5’-ttcgatccgagacatggtacccttgtacagctcgtccatgc-3’ as reverse primers. The red fluorescent variant mApple-GEF-H1 C53R was a kind gift by Perihan Nalbant (University Duisburg-Essen). Briefly, this plasmid was generated via Gibson assembly. The mApple insert was amplified via PCR using the primers 5’-ctcgtttagtgaaccgtcagGCCACCATGGTGAGCAAG-3’ and 5’-ttcgatccgagacatggtaccCTTGTACAGCTCGTCCATG-3’ and the backbone (EGFP-GEF-H1 C53R) was digested with KpnI and EcoR1. The Rho activity sensor delCMV-mCitrine-Rhotekin-GBD was already described (Kamps et al., 2020) and the corresponding control construct delCMV-mCitrine was also constructed previously (Nanda et al., 2023). All ROCK1 and ROCK2 expression vectors and amino acid numbering are based on the canonical human isoforms (UniProt sequence Q13464 and O75116, respectively). CMV promotor driven ROCK1 full length expression constructs mCherry-ROCK1 and mCherry-ROCK2 were constructed via Gateway cloning using the destination vector pDEST-CMV-N-mCherry (Addgene plasmid #123215) and the entry vector R777-E283 Hs.ROCK1 (Addgene plasmid #70567) or R777-E285 Hs.ROCK2 (Addgene plasmid #70569), respectively (Agrotis et al., 2019). The N-terminally truncated ROCK1 constructs were all generated by 1 fragment Gibson assembly using mCherry-ROCK1 as template. In all cases, 5’-ggtgccaacttttttgtac-3’ was used as reverse primer, combined with the following forward primers: mCherry-RBD-PHC1 ROCK1: 5’-tgtacaaaaaagttggcaccatgctaaccaaagatattgaaatattaag-3’, mCherry-dPHC1 ROCK1: 5’-tgtacaaaaaagttggcaccaaaattgatagaaagaaagctaatac-3’, mCherry-mPHC1 ROCK1: 5’-tgtacaaaaaagttggcacctccggttctggaagtggaGAGTCAAGAATTGAAGGTTG-3’. Previous work already established suitable borders for split PH and C1 domains based on the ROCK1 PHC1 tandem domain (Wen et al., 2008). In the current study, the generation of the analogous split PH and C1 domains was based on these previously established sequence borders and carried out via Gibson assembly using mCherry-mPHC1-ROCK1 as the template. For mPH-ROCK1, a 1 fragment Gibson was carried out with the 5’-atttccaaaatcacaaaggcccatgtaaagtaagttatgatg-3’ forward primer (containing an additional 3xGS linker sequence to connect the N- and C-terminal PH domain sequences) (Wen et al., 2008)and 5’-gcctttgtgattttggaaattag-3’ as reverse primer. mCherry-mC1-ROCK1 was generated with a 2 fragment Gibson assembly, both obtained using mCherry-mPHC1-ROCK1 as the template. The insert was generated using 5’- cctccggttctggaagtggacatgagtttattcctacactc-3’ as forward and 5’-acaaattaagtcctctttcttatc-3’ as reverse primer and the backbone was amplified with 5’-agaaagaggacttaatttgttaatcaactttcttgtacaaagtggtg-3’ and 5’-tccacttccagaaccggag-3’. To create mPHC1-ROCK1 RhoA binding point mutants, two double-stranded synthetic DNA fragments were synthesized (Genestrands; Eurofins). Fragment A contained the ROCK1 coding sequence from amino acids 1119-1256 and mutations F1174A-L1199A-E1203K. Fragment B contained the ROCK1 coding sequence from amino acids 1257-1254 and the A1292R mutation. Fragment A was amplified with forward primer 5’-agtcaagaattgaaggttggctttcagtaccaaatagagg-3’ and reverse primer 5’-ctagggcagggggtggcttaaaaacatg-3’. Fragment B was amplified with forward primer 5’-taagccaccccctgccctagagtgtcgaag-3’ and reverse primer 5’-gattaactagtttttccagatgtatttttgaccactttccgg-3’. mCherry-mPHC1-ROCK1 was used as the template for the backbone. To generate mCherry-mPHC1 mutA^PH^, the backbone was amplified using 5’-tctggaaaaactagttaatcaac-3’ as forward and 5’-ctagggcagggggtggcttaaaaacatg-3 as reverse primer and then the Gibson assembly reaction was carried out with the amplified backbone and Fragment B. To generate mCherry-mPHC1 mutB^PH^, the backbone was amplified using 5’-taagccaccccctgccctagagtgtcgaag-3’ as forward and 5’-ccaaccttcaattcttgac-3 as reverse primer and then the Gibson reaction was carried out with the amplified backbone and Fragment A. For mCherry-mPHC1 mutC^PH^, the backbone was amplified using ’5’-tctggaaaaactagttaatcaac-3’ and 5’-ccaaccttcaattcttgac-3’. Then the Gibson reaction was carried out with the amplified backbone, Fragment A and Fragment B. Full length, CMV promotor-driven ROCK1 expression constructs that harbor point mutations in the Rho binding domains were generated via Gibson assembly or using a QuickChange protocol. For mCherry-ROCK1 mutC^PH^ the backbone was amplified from mCherry-ROCK1 using 5’-aacatcacgaAGAGATATGCTGCTGTTAG-3’ as forward and 5’- taacgtgagcCAGTTTATCTATGTCCAATACC-3’ as reverse primer. The insert was amplified from mCherry-mPHC1 mutC^PH^ with forward primer 5’-agataaactgGCTCACGTTAGACCTGTAAC-3’ and reverse primer 5’-gcatatctctTCGTGATGTTACATCATAACTTAC-3’. All three RBD mutant containing plasmids, mCherry-ROCK1RBDmut^RBD^, mCherry-ROCK1mut^RBD^, and mCherry-ROCK1mut^RBD^mutC^PH^, were generated via QuickChange using 5’-cccttaaaacacaggctgttaccacattggc-3’ as forward and 5’-tcgattcattatttctgccaatgtggtaacagcct-3’ as reverse primer, and either mCherry-ROCK1RBD, mCherry-ROCK1 or mCherry-ROCK1mutC^PH^ as templates. The “Molecular Activity Painting” perturbation construct mTurquoise2-eDHFR-GEF-H1 C53R and the dimerizer-presenting artificial receptor construct Dimerizer-PARC-CCL-mCitrine-Halotag to locally stimulate cell contraction signal network activity were already described previously in (Kamps et al., 2020) and (Chen et al., 2002) respectively. The ROCK1 perturbation constructs mTurquoise2-ROCK1-eDHFR^4^ and mTurquoise2-CA-ROCK1-eDFHR were generated via inverse PCR with subsequent Gibson assembly. First, the GEF-H1 C53R insert of mTurquoise2-eDHFR-GEF-H1 C53R was removed via inverse PCR using primers 5’-TAACTCAATTGTTGTTGTTAACTTGTTTATTGC-3’ and 5’-CCGCCGCTCCAGAATCTC-3’ followed by blunt end ligation of the PCR product. The resulting plasmid was used as a backbone for Gibson assembly after restriction digestion using XhoI (NEB). The inserts for Gibson assembly were amplified from mCherry-ROCK1 (amino acids 1-1354 for mTurqoise-ROCK1-eDHFR and 1-477 for mTurqoise2-CA-ROCK1-eDHFR) with 5’-ctggactcgtacaagatatcaatgtcgactggggacagttttg-3’ as forward and 5’-gcaatcagactgatcatagccgatccacttccagaaccggaactagtttttccagatgtatttttgacc-3’ as reverse primer (for mTurquoise2-ROCK1-eDHFR), or 5’-gatctggactcgtacaagatatcatccggttctggaagtggatcgactggggacagttttg-3’ as forward and 5’-gccgcaatcagactgatcatagctccacttccagaaccggatagatttcttctttgatttccctcttc-3’ as reverse primer (for mTurquoise2-CA-ROCK1-eDFHR). The mutated perturbation construct mTurquoise2-ROCK1Δ^PHC1^-eDHFR (vector ID: VB250121-1429rnx) was constructed by the company VectorBuilder (Chicago, USA) starting with the mTurquoise2-eDHFR and mCherry-ROCK1 constructs described above. The mutated perturbation construct mTurquoise2-ROCK1 mutC^PH^-eDHFR was generated via Gibson assembly: The backbone was amplified from mTurquoise2-ROCK1-eDHFR with 5’-aacatcacgaAGAGATATGCTGCTGTTAG-3’ as forward and 5’-taacgtgagcCAGTTTATCTATGTCCAATACC-3’ as reverse primer and the insert was amplified from mCherry-mPHC1 mutC^PH^ with forward primer 5’- agataaactgGCTCACGTTAGACCTGTAAC-3’ and reverse primer 5’-gcatatctctTCGTGATGTTACATCATAACTTAC-3’. The expression vector mCherry-NMHC2a encoding non-muscle Myosin heavy chain IIa (*MYH9*, human), was already described previously (Kowalczyk et al., 2022).

### Microscopy

TIRF microscopy was performed on an Olympus IX-81 microscope, equipped with an Apo TIRF 60 × 1.45 NA oil immersion objective, a TIRF-MITICO motorized TIRF illumination combiner and a ZDC autofocus device. Four different lasers were employed for imaging cells: 514 nm OBIS diode laser (150 mW) (Coherent, Inc., Santa Clara, USA) and Cell R diode lasers (Olympus) with wavelength 405 nm (50 mW), 445 nm (50 mW) and 561 nm (100 mW). In addition, wide-field illumination was performed using the Spectra X light engine (Lumencor). Signals were detected using an EMCCD camera (C9100-13; Hamamatsu, Herrsching am Ammersee, Germany) at medium gain without binning. Live-cell imaging was carried out at 33 °C in CO--independent HEPES-stabilized imaging medium (Pan Biotech Cat.no.: P04-01163) supplemented with 10% FBS and at the indicated frame rates.

### Functionalized Molecular Activity Painting

The original molecular activity painting method was described previously (Chen et al., 2017), and a simplified variant used in this work was developed subsequently (Kamps et al., 2020). Briefly, 1.0-1.4x10^5^ U2OS cells were plated into each well of a 6-well plate (Cell+ Sarstedt Ref.: 83.3920.300) and transfected on day 2 with a dimerizer-presenting artificial receptor construct (dimerizer-PARC), a construct containing *E. coli* dihydrofolate reductase (eDHFR) fused to a protein of interest (POI), and fluorescent-labelled readout protein(s). On the next day, the cells were detached by incubation with 10 mM EDTA (in DPBS, pH 7.4) for 25-30 min without trypsin or by incubation with the Enzyme Free Cell Dissociation solution (EMD Millipore Corp, S-014-B) for 10 min. The cells were washed 2x (EDTA-detachment) or 1x (Enzyme Free Dissoziation Solution-detachment) with serum-free medium followed by centrifugation. After washing, cells were resuspended in 200 µl serum-free medium and plated onto the functionalized glass-bottom dishes (see next paragraph). After incubation at 37 °C, 5% CO-for 30 min, 200 µl 20% FBS-containing medium was added and incubation continued overnight.

On day 2, glass-bottom dishes (8-well LabTek I, VWR) are coated with 0.1% poly-L-lysine hydrobromide (Sigma-Aldrich) at 4°C overnight. Next day, the wells were washed 3x with DPBS followed by biotinylation of adsorbed poly-L-lysine using 1 mg/ml EZ-link Sulfo-NHS-Biotin (Fisher Scientific) in DPBS for 1 hr at room temperature. During incubation, two solutions were prepared on ice (volume for 3 wells): a) streptavidin-solution: 2 µl Streptavidin (Serva, 1 mg/ml) + 1 µl streptavidin Alexa Fluor 730 (Life Technologies, 0.1 mg/ml) + 12 µl DPBS and b) antibody solution: 3 μl biotin-labeled anti-VSVG-antibody (Abcam ab34774, 1 mg/ml) + 12 μl DPBS. After 3 washes with DPBS, all remaining DPBS was aspirated from the biotinylated glass surface. The two prepared solutions were quickly mixed and 5x 2 µl drops were added on the dry glass surface of each well to finalize functionalization by streptavidin- and antibody addition in the resulting small circular patterns (diameter: 1-2 mm). Following a minimum incubation period of 2 min, the wells were washed 3x with DPBS and then stored at 4 °C, submerged in DPBS, for up to 45 min prior to cell seeding. The fluorescence signal of streptavidin Alexa Fluor 750 enabled the identification of functionalized regions during imaging.

Covalent labelling for chemo-optogenetic perturbation was carried out by incubating with 20 μM of the Nvoc-TMP-Cl photocaged chemical dimerizer (Chen et al., 2017) in HEPES-stabilized imaging medium for 60 minutes. Excess dimerizer was removed by three consecutive washes with HEPES-stabilized imaging medium, followed by a fourth wash after an additional 30-minute incubation. Photo uncaging was achieved using a 405 nm laser focused to a spot through the FRAP mode of the TIR-MITICO motorized TIRF illumination combiner. The process was carried out for 400 ms at 180 nW (20% laser power), with the laser power measured at the 60x objective. Since the perturbation is irreversible, only a single light pulse was applied to each cell.

### Image and video analysis

All image analysis was conducted using ImageJ (http://imagej.nih.gov/ij/), and figures were created with Adobe Illustrator. Kymograph analysis was performed using the multi kymograph plugin in ImageJ. Cell movies were stabilized using the ImageJ plugins Image Stabilizer and Image Stabilizer Log Applier (https://imagej.net/plugins/image-stabilizer). Statistical analyses and generation of plots were carried out with Prism (GraphPad). All statistical tests were two-sided.

### Analysis of perturbation response kinetics

The kinetics of Rho, GEF-H1, ROCK1 or Myosin fluorescence signal changes following chemo-optogenetic perturbations were measured at the local perturbation site, as described previously (Chen et al., 2017). The fluorescence signal profiles for the perturbation and response, shown in Figure 3 and 5, are normalized to the time point right before perturbation.

### Analysis of cell contraction signal network dynamics

As an unbiased analysis of signal network dynamics, we employed a custom-made ImageJ macro as previously described (Graessl et al., 2017) to determine the standard deviation, pulse frequency, and pulse amplitude of proteins that are dynamically recruited to the plasma membrane with minor modifications.

### AlphaFold modelling

The model of human ROCK1 kina se (UNIPROT Q13464) was created by predicting dimers of the fragments 1-600, 500-700, 600-800, 500-1000, and 900-1354 using AlphaFold Multimer version 2.2.0 (Evans et al., 2021) and joining them together using coot (Emsley and Cowtan, 2004), followed by geometry relaxation with alpha-helical constraints. For the complex of RhoA with the ROCK PH domain, fragments 1120-1334 of ROCK1, 1152-1374 of ROCK2 (UNIPROT O75115) and 950-1173 of let-502 from C. elegans (UNIPROT P92199) were predicted using AlphaFold3^36^ in complex with human or *C. elegans* RhoA (UNIPROT P61586 or Q22038) (Figure S5A). The PAE and LDDT scores were very significant for all three complexes. The PAE plot reveals a stronger connection between the C1 domain and RhoA for the let-502 compared to the human PHC1 domains, indicating that the main binding interface for the human proteins is the PH domain, although some residues of the C1 domain might also contribute (Figure S5B). The RhoA-PHC1 interface in the *C. elegans* proteins has therefore a slightly different residue conservation pattern. This is also visible in the ipSAE plots (Dunbrack, 2025), where the best C. elegans model shows again higher scores for the C1 domain (Figure S5C). Corroborating this observation, the different models for the *C. elegans* protein predict the C1 domain in quite similar positions, whereas the relative orientation of the C1 domain compared to the PH and RhoA domains varies. Still, there are some residues with good PAE scores relative to RhoA, and the steric positioning of RhoA by the C1 domain might also contribute to the overall affinity. For the construct mPH-ROCK1 where the two sections of the PH domain are connected by a long linker, AF3 predicts the same complex as for the construct that includes the ROCK1-C1 domain, but with lower ipTM scores (0.75 vs 0.25), indicating that the C1 domain indeed contributes to the stability of the complex.

The putative membrane-bound model (Figure 6F) was built by superimposing an AlphaFold2 prediction of human ROCK1 residues 900-1354, the crystal structure of human ROCK1 residues 946-1015 (PDB ID 1S1C), and predictions of ROCK1 900-1354 in complex with RhoA. The PHC1-RhoA domains were oriented manually so that the RhoA C-termini and the positive charges of the PHC1 domains face the putative plasma membrane surface, which is easily achieved due to the disordered linker formed by residues 1103-1121 between the coiled-coil and the start of the PH domain of ROCK1.

## Supporting information

Movie_Figure1_top

Movie_Figure1_middle

Movie_Figure1_bottom

Movie_Figure2B-D_top

Movie_Figure2B-D_middle

Movie_Figure2B-D_bottom

Movie_Figure2I-K_top

Movie_Figure2I-K_middle

Movie_Figure2I-K_bottom

Movie_Figure3D-F_top

Movie_Figure3D-F_middle

Movie_Figure3D-F_bottom

Movie_Figure3J-K_top

Movie_Figure3J-K_bottom

Movie_Figure4A-C_top

Movie_Figure4A-C_bottom

Movie_Figure4F-G_top

Movie_Figure4F-G_middle

Movie_Figure4F-G_bottom

Movie_Figure5B-C_top

Movie_Figure5B-C_middle1

Movie_Figure5B-C_middle2

Movie_Figure5B-C_bottom

Movie_FigureS3_top

Movie_FigureS3_bottom

## Acknowledgements

We want to thank Dominic Kamps (TU Dortmund) for generating the mTurquoise2-NES-Linker-eDHFR construct that was used for further plasmid construction and the lab of Perihan Nalbant (University Duisburg-Essen) for generating the mApple-GEF-H1 C53R plasmid that was used to confirm the network dynamics of Rho in comparison to GEF-H1 C53R. We are thankful for fruitful discussions with Perihan Nalbant, Peter Bieling (MPI Dortmund) and Rudolf Merkel (FZ Jülich). We would also like to thank Ricarda Lüttig (TU Dortmund), Arya Sachan (TU Dortmund) and Lejla Maksumic (TU Dortmund) for scientific input. This work was supported by the Deutsche Forschungsgesellschaft DFG Heisenberg Programme grant DE 823/8-1, the DFG project grant DE 823/9-1 and the DFG principal investigator grant DE 823/10-1 to L.D.

## Author contributions

L.D. and C.G. designed the research. C.G. performed and analyzed all experiments. L.D. supervised all experiments. C.G. and L.D. wrote the manuscript. I.R.V. and L.D. performed AlphaFold predictions and structure related analyses.

## Supplement

**Figure S1 (related to Introduction and Figure1):**
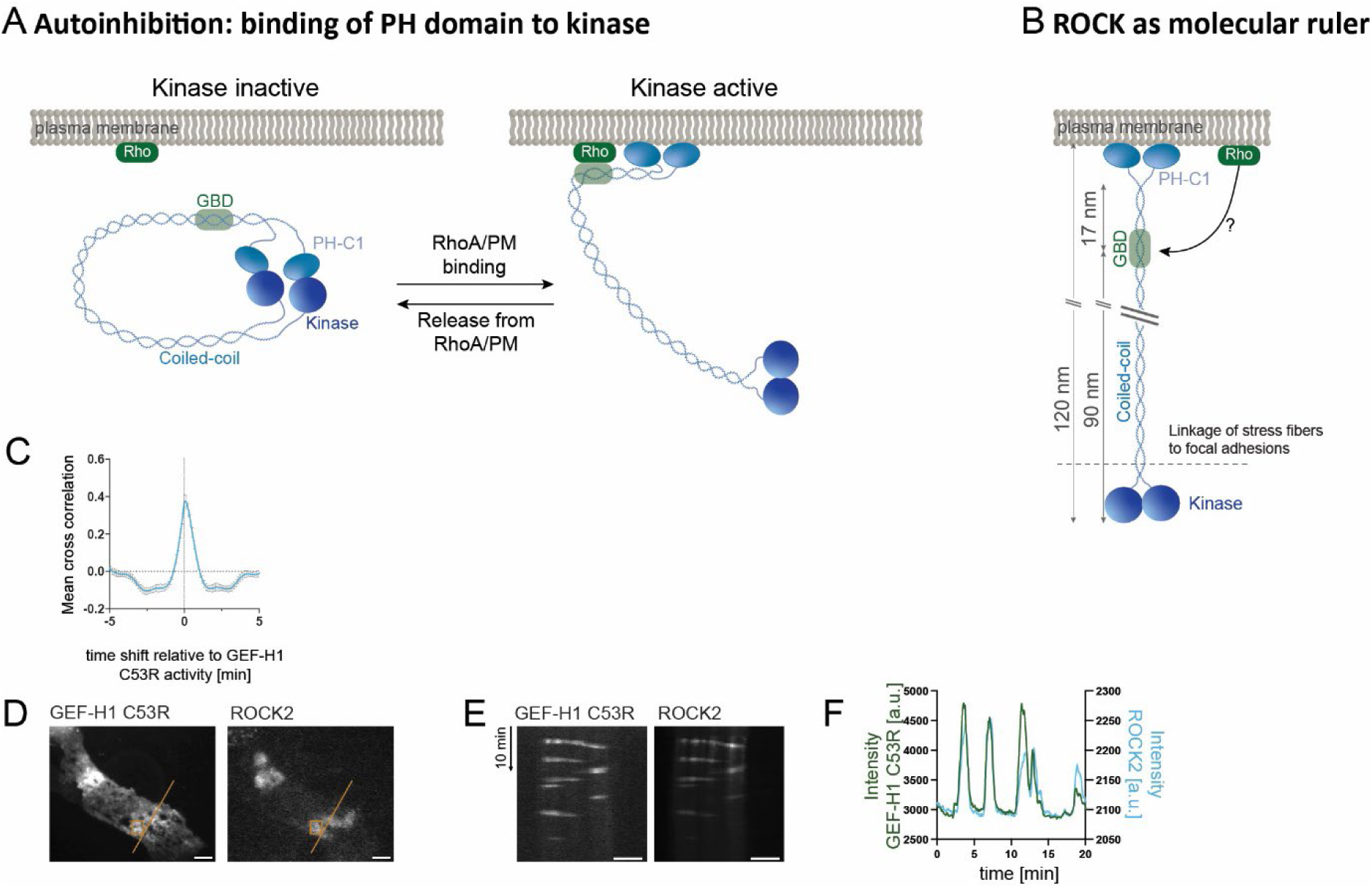
**A** Classical model of ROCK regulation via autoinhibition, in which the PHC1 domain is thought to inhibit the kinase domain. **B** More recent model in which the coiled-coil domain positions the kinase domain at a defined distance from the plasma membrane. In this model, ROCK1 is thought to be constitutively bound to the plasma membrane. How RhoA would activate ROCK in this conformation is unclear. **C** Cross correlation analysis of GEF-H1 and ROCK1 (n=59 cells, 4 independent experiments, see Methods for details). **D**-**F** Analysis of ROCK2 plasma membrane recruitment dynamics. **D** TIRF images of a representative cell co-expressing mCitrine-GEF-H1 C53R and mCherry-ROCK2 full length. **E** Kymograph corresponding to the orange lines in **D**. **F** Kinetics of fluorescence signals in rectangular regions of interest in **D**. Error bars represent standard error of the mean, scale bar = 10 µm.

**Figure S2 (related to Figure 3):**
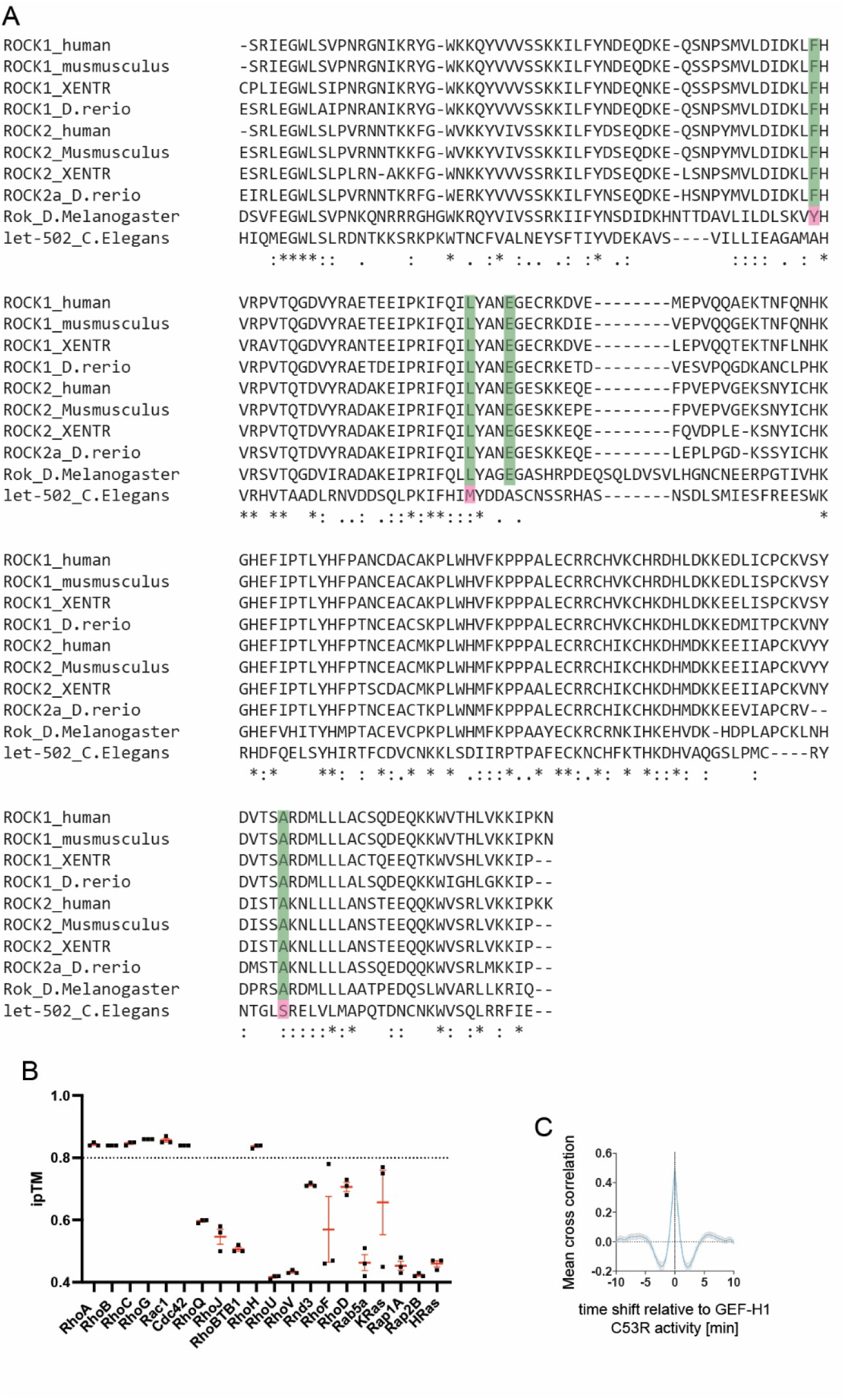
**A** Protein Sequence alignment of the ROCK1 and analogous PH domain using *Clustal 2.1 multiple sequence alignment* web tool. **B** The interface predicted template modelling score (ipTM) of various GTP bound small GTPases in complex with the PHC1 tandem domain obtained via Alphafold3 predictions. The dotted line represents the 0.8 threshold that typically discriminates true vs false positives. **C** Cross correlation analysis of GEF-H1 and mPHC1-ROCK1. Error bars represent standard error of the mean from 3 independent predictions or experiments.

**Figure S3 (related to Figure 4):**
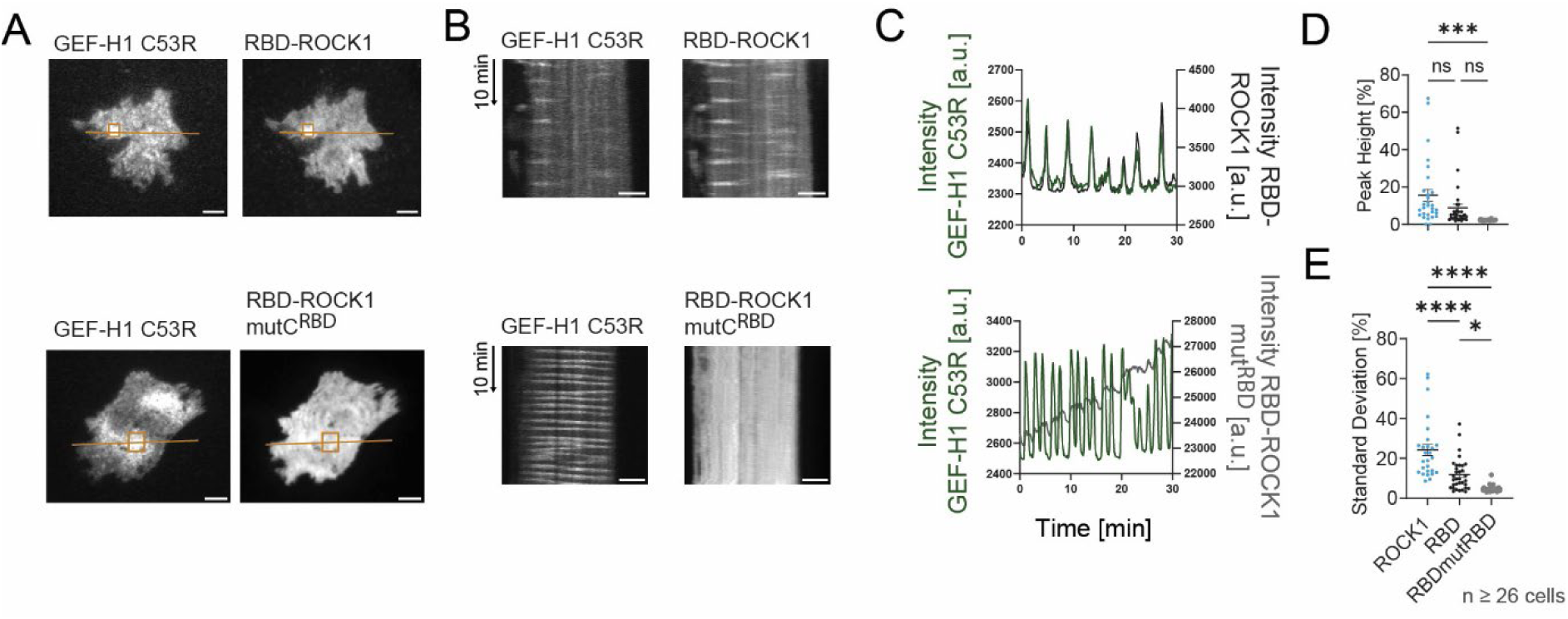
**A** TIRF images of representative cells co-expressing mCitrine-GEF-H1 and mCherry-RBD-ROCK1 (top) or mCherry-RBD-ROCK1 mut^RBD^ (bottom). **B** Kymographs corresponding to lines in **A**. **C** Kinetics of fluorescence signals in rectangular regions of interest in **A**. **D**-**E** Quantitative analysis of plasma membrane recruitment dynamics of mCherry-RBD-ROCK1 signals shown in **A**-**C.** The control data (ROCK1 full length) is the same as shown in Figure 4, included here again to allow comparison with the treatment group. **D** Quantification of pulse height. **E** Quantification of the local standard deviation over time (3 independent experiments). Error bars represent standard error of the mean, scale bar: 10 µm.

**Figure S4 (related to Figure 6):**
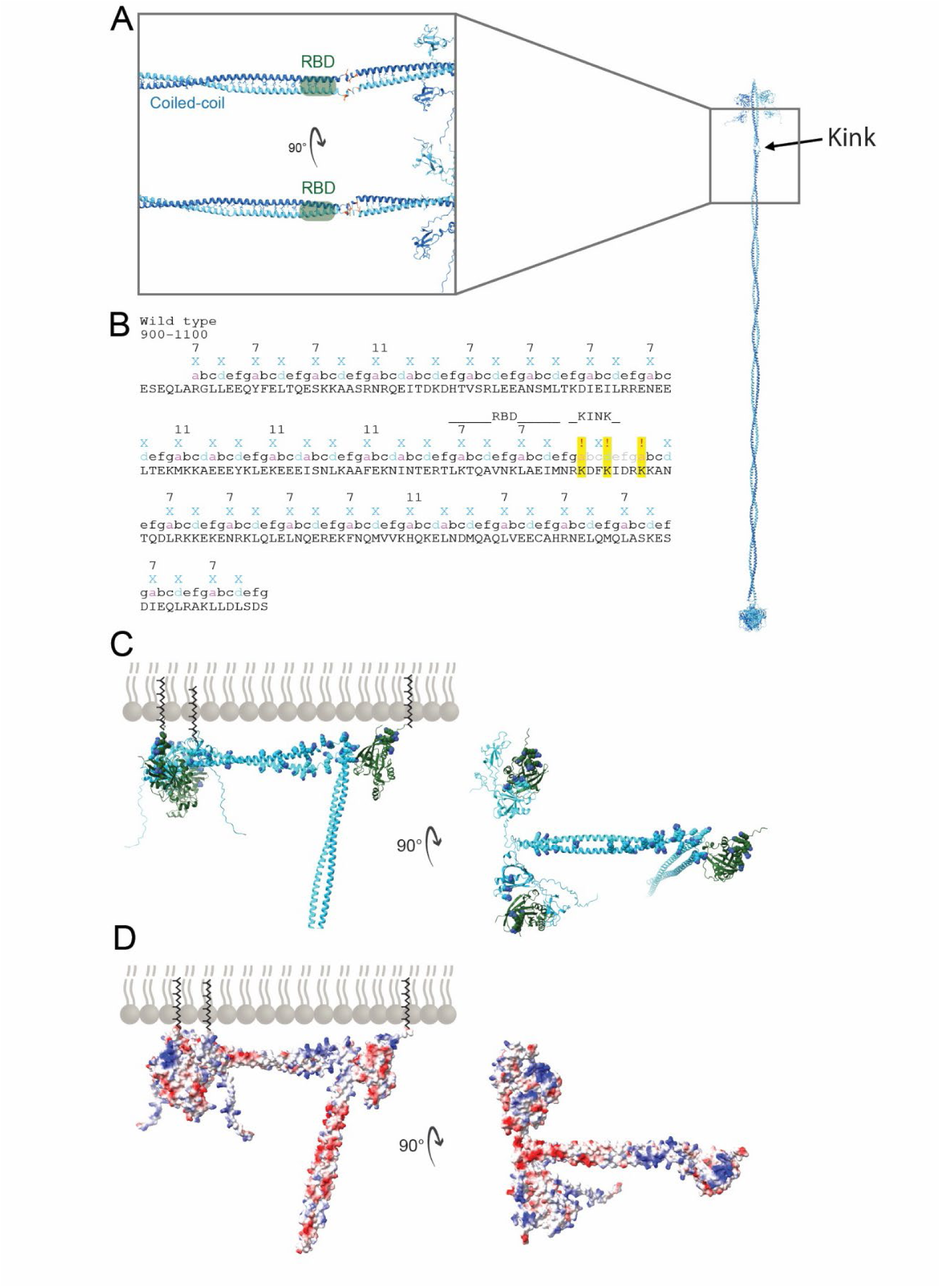
AlphaFold prediction of a kink in the ROCK1 coiled-coil domain. **A** Zoom into a region of a full length ROCK1 Alphafold2 prediction. The full-length AlphaFold structure is based on a combination of multiple predictions of successive ROCK1 fragments (see AlphaFold methods). The predicted kink C-terminal to the RBD (Rho-binding domain) is due to a disruption in the repetitive coiled-coil amino sequence motif. **B**: Amino acid sequence of the coiled coil and kink regions of ROCK1 (corresponding to AA 900-1100). ‘X’ marks positions, in which the amino acids point towards the core of the coiled-coil structure in the Alphafold2 prediction. Note that the majority of these amino acids have hydrophobic character. ‘abcdefg’ represents a single, canonical coiled-coil heptad repeat, ‘abcdabcdefg’ represents a single, non-canonical hendecad repeat. The numbers 7 or 11 highlight the start of heptad or hendecad repeats, respectively. Positions highlighted in yellow and with an exclamation mark (‘!’) indicate amino acids that would be expected to be hydrophobic based on canonical coiled-coil heptad repeats, but instead are hydrophilic and face away from the coiled-coil core in the Alphafold2 prediction. **C** Alphafold2 prediction of the RBD and PHC1 domains in complex with three active Rho molecules. A preferred perpendicular orientation of the ROCK1 relative to the plasma membrane due to interactions with three membrane bound Rho molecules is indicated, which requires the predicted kink shown above. The electrostatic surface of ROCK1 (right) reveals a cluster of positively charged amino acids at the predicted membrane interaction surface. Rotating the protein by 90° reveals the full plasma membrane interaction surface and highlights the high amount of positive charges (dark blue) in this region. The predicted interaction between active Rho and the RBD region is similar to the published crystal structure (PDB ID 1S1C), which was derived from a complex of Rho with a ROCK1 residues 946-1015, comprising the RBD region of the coiled-coil, an upstream region of the coiled-coil, but lacking the majority of amino acids downstream of the RBD binding site. **D** Electrostatic surface of the model shown in C.

**Figure S5 (related to Methods):**
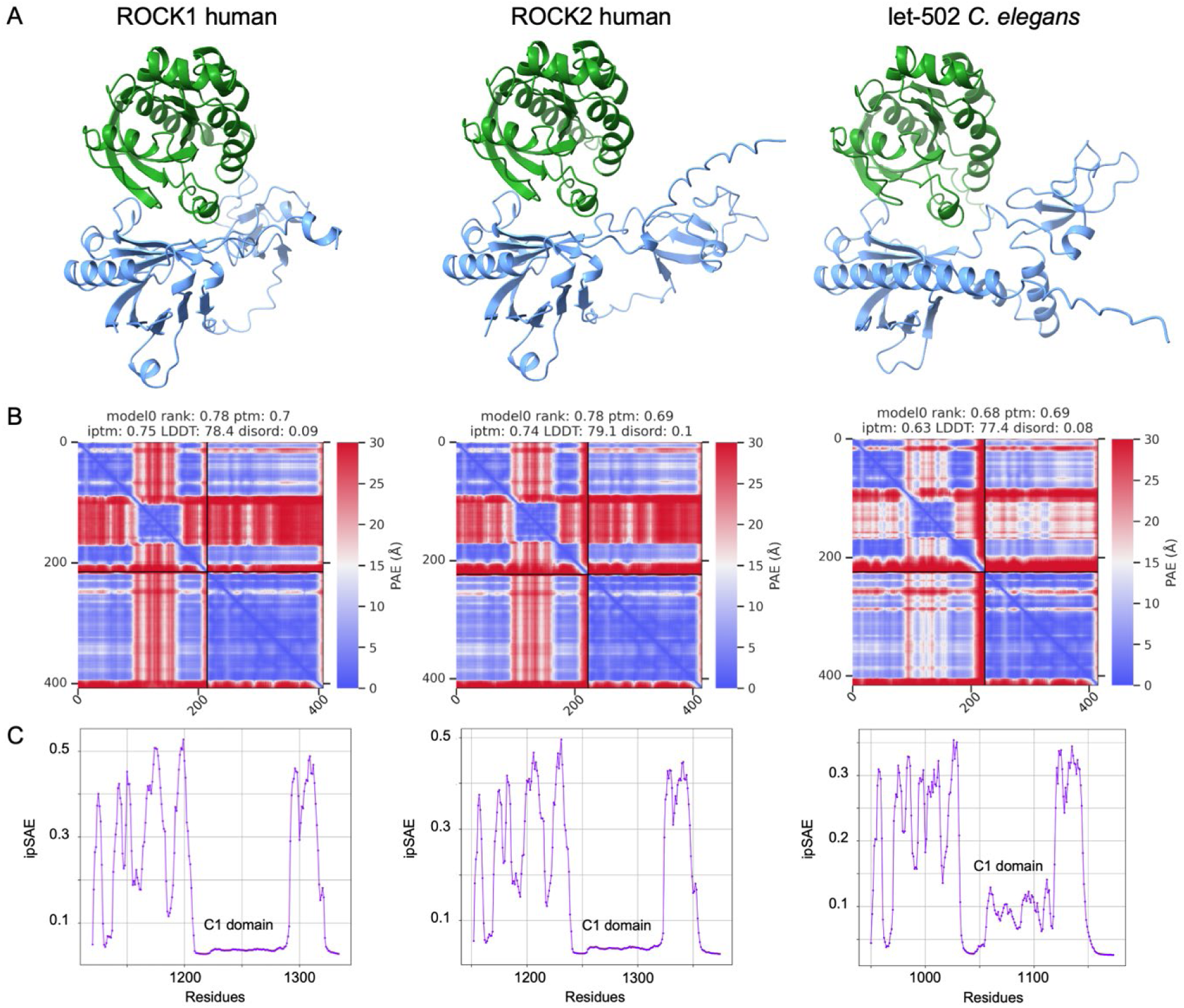
AlphaFold predictions of the complex of RhoA with ROCK PH domains. **A** Predicted structures of the ROCK PH domain (blue) in complex with human or C. elegans RhoA (green), using AlphaFold3 (Abramson et al., 2024). **B** PAE plots for the complexes shown in **A**. **C** ipSAE plots of the predictions shown in **A**.

## Movie Legends

Movie_Figure1_top: GEF-H1 C53R expression stimulates cell contraction network dynamics to form pulse- and wave-like activity patterns. Time-lapse TIRF videos of mCitrine-GEF-H1 C53R and mCherry in a representative U2OS cell. Imaging started 42 h after plating on collagen-coated glass-bottom dishes. Images were collected with a frame rate of 7.5/min. Scale bars, 10 µm.

Movie_Figure1_middle: Rho activity strongly correlates with GEF-H1 C53R stimulated cell contraction signal network activity in space and time. Time-lapse TIRF videos of mApple-GEF-H1 C53R and the Rho activity sensor mCitrine-Rhotekin-GBD in a representative U2OS cell. Imaging started 42 h after plating on collagen-coated glass-bottom dishes. Images were collected with a frame rate of 7.5/min. Scale bars, 10 µm.

Movie_Figure1_bottom and Figure2B-D_top: ROCK1 strongly correlates with GEF-H1 C53R stimulated cell contraction signal network activity at the plasma membrane. Time-lapse TIRF videos of mCitrine-GEF-H1 C53R and mCherry-ROCK1 full length in a representative U2OS cell. Imaging started 42 h after plating on collagen-coated glass-bottom dishes. Images were collected with a frame rate of 7.5/min. Scale bars, 10 µm.

Movie_Figure2B-D_top: ROCK1 is strongly recruited to GEF-H1 C53R stimulated cell contraction signal network activity at the plasma membrane. Time-lapse TIRF videos of mCitrine-GEF-H1 C53R and mCherry-ROCK1 full length in a representative U2OS cell. Imaging started 42 h after plating on collagen-coated glass-bottom dishes. Images were collected with a frame rate of 7.5/min. Scale bars, 10 µm.

Movie_Figure2B-D_middle: N-terminal truncated RBD-PH-ROCK1 is strongly recruited to GEF-H1 C53R stimulated cell contraction signal network activity at the plasma membrane. Time-lapse TIRF videos of mCitrine-GEF-H1 C53R and mCherry-RBD-PHROCK1 in a representative U2OS cell. Imaging started 42 h after plating on collagen-coated glass-bottom dishes. Images were collected with a frame rate of 7.5/min. Scale bars, 10 µm.

Movie_Figure2B-D_bottom: N-terminal truncated PHC1-ROCK1 is strongly recruited to GEF-H1 C53R stimulated cell contraction signal network activity at the plasma membrane. PHC1-ROCK1 does not contain any of the previously identified GTPase-binding domains of ROCK1. Time-lapse TIRF videos of mCitrine-GEF-H1 C53R and mCherry-dPHC1-ROCK1 in a representative U2OS cell. Imaging started 42 h after plating on collagen-coated glass-bottom dishes. Images were collected with a frame rate of 7.5/min. Scale bars, 10 µm.

Movie_Figure2I-K_top: The monomeric PHC1 tandem domain of ROCK1 (mPHC1-ROCK1) is recruited to GEF-H1 C53R stimulated cell contraction signal network activity at the plasma membrane. mPHC1-ROCK1 does not contain any of the previously identified GTPase-binding domains of ROCK1 nor any known dimerization motifs. Time-lapse TIRF videos of mCitrine-GEF-H1 C53R and mCherry-mPHC1-ROCK1 in a representative U2OS cell. Imaging started 42 h after plating on collagen-coated glass-bottom dishes. Images were collected with a frame rate of 7.5/min. Scale bars, 10 µm.

Movie_Figure2I-K_middle: mPH-ROCK1 (PH part of PHC1 domain) is not efficiently recruited to GEF-H1 C53R stimulated cell contraction signal network activity at the plasma membrane. Time-lapse TIRF videos of mCitrine-GEF-H1 C53R and mCherry-mPH-ROCK1 in a representative U2OS cell. Imaging started 42 h after plating on collagen-coated glass-bottom dishes. Images were collected with a frame rate of 7.5/min. Scale bars, 10 µm.

Movie_Figure2I-K_bottom: mC1-ROCK1 (C1 part of PHC1 domain) is not efficiently recruited to GEF-H1 C53R stimulated cell contraction signal network activity at the plasma membrane. Time-lapse TIRF videos of mCitrine-GEF-H1 C53R and mCherry-mPH-ROCK1 in a representative U2OS cell. Imaging started 42 h after plating on collagen-coated glass-bottom dishes. Images were collected with a frame rate of 7.5/min. Scale bars, 10 µm.

Movie_Figure3D-F_top: The mutated monomeric PHC1 tandem domain of ROCK1 (mPHC1-ROCK1 mutA^PH^) with the A1292R point mutation is not efficiently recruited to GEF-H1 C53R stimulated cell contraction signal network activity at the plasma membrane. Time-lapse TIRF videos of mCitrine-GEF-H1 C53R and mCherry-mPH-ROCK1 in a representative U2OS cell. Imaging started 42 h after plating on collagen-coated glass-bottom dishes. Images were collected with a frame rate of 7.5/min. Scale bars, 10 µm.

Movie_Figure3D-F_middle: The mutated monomeric PHC1 tandem domain of ROCK1 (mPHC1-ROCK1 mutB^PH^) with the L1199A/F1174A/E1203K point mutations is not efficiently recruited to GEF-H1 C53R stimulated cell contraction signal network activity at the plasma membrane. Time-lapse TIRF videos of mCitrine-GEF-H1 C53R and mCherry-mPH-ROCK1 in a representative U2OS cell. Imaging started 42 h after plating on collagen-coated glass-bottom dishes. Images were collected with a frame rate of 7.5/min. Scale bars, 10 µm.

Movie_Figure3D-F_bottom: The mutated monomeric PHC1 tandem domain of ROCK1 (mPHC1-ROCK1 mutC^PH^) with the L1199A/F1174A/E1203K/A1292R point mutations is not recruited to GEF-H1 C53R stimulated cell contraction signal network activity at the plasma membrane. Time-lapse TIRF videos of mCitrine-GEF-H1 C53R and mCherry-mPH-ROCK1 in a representative U2OS cell. Imaging started 42 h after plating on collagen-coated glass-bottom dishes. Images were collected with a frame rate of 7.5/min. Scale bars, 10 µm.

Movie_Figure3J-K_top: Rapid chemo-optogenetic plasma membrane recruitment of GEF-H1 C53R leads to recruitment of the monomeric PHC1 domain of ROCK1 at a distinct site at the plasma membrane. TIRF video-microscopy of a representative U2OS cell, expressing the perturbation construct mTurquoise2-eDHFR-GEF-H1C53R (left), the response construct mCherry-mPHC1-ROCK1 (right) and the artificial receptor mCitrine-VSVG HaloTag-PARC. A focused light pulse at 405nm was applied at timepoint 0 to target the perturbation construct to a small region of the plasma membrane. Images were collected with a frame rate of 6/min. Scale bars, 10 µm.

Movie_Figure3J-K_bottom: Rapid chemo-optogenetic plasma membrane recruitment of GEF-H1 C53R does not lead to plasma membrane recruitment of the mutated monomeric mPHC1 mutC^PH^ domain of ROCK1 with the L1199A/F1174A/E1203K/A1292R point mutations. TIRF video-microscopy of a representative U2OS cell, expressing the perturbation construct mTurquoise2-eDHFR-GEF-H1C53R (left), the response construct mCherry-mPHC1mutC^PH^-ROCK1 (right) and the artificial receptor mCitrine-VSVG HaloTag-PARC. A focused light pulse at 405nm was applied at timepoint 0 to target the perturbation construct to a small region of the plasma membrane. Images were collected with a frame rate of 6/min. Scale bars, 10 µm.

Movie_Figure4A-C_top: Full length ROCK1 is strongly recruited to GEF-H1 C53R stimulated cell contraction signal network activity at the plasma membrane. Time-lapse TIRF videos of mCitrine-GEF-H1 C53R and mCherry-ROCK1 in a representative U2OS cell. Imaging started 42 h after plating on collagen-coated glass-bottom dishes. Images were collected with a frame rate of 7.5/min. Scale bars, 10 µm.

Movie_Figure4A-C_bottom: Reduced plasma membrane recruitment of mutated full length ROCK1 mutC^PH^ with the L1199A/F1174A/E1203K/A1292R point mutations to GEF-H1 C53R stimulated cell contraction signal network activity at the plasma membrane. Time-lapse TIRF videos of mCitrine-GEF-H1 C53R and mCherry-ROCK1mutC^PH^ in a representative U2OS cell. Imaging started 42 h after plating on collagen-coated glass-bottom dishes. Images were collected with a frame rate of 7.5/min. Scale bars, 10 µm.

Movie_Figure4F-H_top: Full length ROCK1 is strongly recruited to cell contraction network activity at the plasma membrane. Time-lapse TIRF videos of mCitrine-GEF-H1 C53R and mCherry-ROCK1 in a representative U2OS cell. Imaging started 42 h after plating on collagen-coated glass-bottom dishes. Images were collected with a frame rate of 7.5/min. Scale bars, 10 µm.

Movie_Figure4F-H_middle: Reduced plasma membrane recruitment of mutated full length ROCK1 mutC^RBD^ with the N1004T/L1005T point mutations to cell contraction network activity at the plasma membrane. Time-lapse TIRF videos of mCitrine-GEF-H1 C53R and mCherry-ROCK1mut^RBD^ in a representative U2OS cell. Imaging started 42 h after plating on collagen-coated glass-bottom dishes. Images were collected with a frame rate of 7.5/min. Scale bars, 10 µm.

Movie_Figure4F-H_bottom: Plasma membrane recruitment of mutated full length ROCK1 mutC^RBD^mutC^PH^ with the L1199A/F1174A/E1203K/A1292R/N1004T/L1005T point mutations to cell contraction network activity at the plasma membrane is further reduced. Time-lapse TIRF videos of mCitrine-GEF-H1 C53R and mCherry-ROCK1mut^RBD^mutC^PH^ in a representative U2OS cell. Imaging started 42 h after plating on collagen-coated glass-bottom dishes. Images were collected with a frame rate of 7.5/min. Scale bars, 10 µm.

Movie_FigureS3A-B_top: Plasma membrane recruitment of the ROCK1 RBD region to cell contraction network activity at the plasma membrane. Time-lapse TIRF videos of mCitrine-GEF-H1 C53R and mCherry-RBD-ROCK1 in a representative U2OS cell. Imaging started 42 h after plating on collagen-coated glass-bottom dishes. Images were collected with a frame rate of 7.5/min. Scale bars, 10 µm.

Movie_FigureS3A-B_bottom: Plasma membrane recruitment of the mutated ROCK1 RBD region with the N1004T/L1005T point mutations to cell contraction network activity is not detectable. Time-lapse TIRF videos of mCitrine-GEF-H1 C53R and mCherry-RBD-ROCK1-mutC^RBD^ in a representative U2OS cell. Imaging started 42 h after plating on collagen-coated glass-bottom dishes. Images were collected with a frame rate of 7.5/min. Scale bars, 10 µm.

Movie_Figure5B-C_top: Rapid chemo-optogenetic plasma membrane recruitment of full length ROCK1 leads to plasma membrane recruitment and activation of Myosin IIa. TIRF video-microscopy of a representative U2OS cell, expressing the perturbation construct mTurquoise2-ROCK1-eDHFR (left), the response construct mCherry-non muscle Myosin II heavy chain IIa (right) and the artificial receptor mCitrine-VSVG HaloTag-PARC. A focused light pulse at 405nm was applied at timepoint 0 to target the perturbation construct to a small region of the plasma membrane. Images were collected with a frame rate of 6/min. Scale bars, 10 µm.

Movie_Figure5B-C_middle1: Rapid chemo-optogenetic plasma membrane recruitment of the C-terminal truncation mutant CA-ROCK1 does not lead to plasma membrane recruitment and activation of Myosin IIa. TIRF video-microscopy of a representative U2OS cell, expressing the perturbation construct mTurquoise2-CA-ROCK1-eDHFR (left), the response construct mCherry-non muscle Myosin II heavy chain IIa (right) and the artificial receptor mCitrine-VSVG HaloTag-PARC. A focused light pulse at 405nm was applied at timepoint 0 to target the perturbation construct to a small region of the plasma membrane. Images were collected with a frame rate of 6/min. Scale bars, 10 µm.

Movie_Figure5B-C_middle2: Rapid chemo-optogenetic plasma membrane recruitment of the C-terminal truncation mutant ROCK1Δ^PHC1^ does not lead to plasma membrane recruitment and activation of Myosin IIa. TIRF video-microscopy of a representative U2OS cell, expressing the perturbation construct mTurquoise2-ROCK1Δ^PHC1^-eDHFR (left), the response construct mCherry-non muscle Myosin II heavy chain IIa (right) and the artificial receptor mCitrine-VSVG HaloTag-PARC. A focused light pulse at 405nm was applied at timepoint 0 to target the perturbation construct to a small region of the plasma membrane. Images were collected with a frame rate of 6/min. Scale bars, 10 µm.

Movie_Figure5B-C_bottom: Rapid chemo-optogenetic plasma membrane recruitment of mutated full length ROCK1mutC^PH^ leads to increased plasma membrane recruitment and activation of Myosin IIa. TIRF video-microscopy of a representative U2OS cell, expressing the perturbation construct mTurquoise2-ROCK1mutC^PH^-eDHFR (left), the response construct mCherry-non muscle Myosin II heavy chain IIa (right) and the artificial receptor mCitrine-VSVG HaloTag-PARC. A focused light pulse at 405nm was applied at timepoint 0 to target the perturbation construct to a small region of the plasma membrane. Images were collected with a frame rate of 6/min. Scale bars, 10 µm.

## Footnotes

1 A companion study by Gierse et al. 2025 that is submitted together with this work, characterized this effect in more detail.

2 A companion study by Gierse et al. 2025 that is submitted together with this work, characterized this effect in more detail.

3 This construct was primarily developed and generated in a companion study by Gierse et al 2025 that is submitted together with this work. The experimental details for the construction of this construct are briefly stated in both papers. For future derivative work that use CMV-mCitrine-GEF-H1 C53R, the companion study should be cited.

4 This construct was primarily developed and generated in a companion study by Gierse et al 2025 that is submitted together with this work. The experimental details for the construction of this construct are briefly stated in both papers. For future derivative work that use mTurquoise2-ROCK1-eDHFR, the companion study should be cited.

## Notes

### Competing Interest Statement

The authors have declared no competing interest.

